# The Golgi Glycoprotein MGAT4D is an Intrinsic Protector of Testicular Germ Cells From Mild Heat Stress

**DOI:** 10.1101/818195

**Authors:** Ayodele Akintayo, Meng Liang, Boris Bartholdy, Frank Batista, Jennifer Aguilan, Jillian Prendergast, Subha Sundaram, Pamela Stanley

## Abstract

Male germ cells are sensitive to heat stress and testes must be maintained outside the body for optimal fertility. However, no germ cell intrinsic mechanism that protects from heat has been reported. Here, we identify the germ cell specific Golgi glycoprotein MGAT4D as a protector of male germ cells from heat stress. *Mgat4d* is highly expressed in spermatocytes and spermatids. Unexpectedly, when the *Mgat4d* gene was inactivated globally or conditionally in spermatogonia, or mis-expressed in spermatogonia, spermatocytes or spermatids, neither spermatogenesis nor fertility were affected. On the other hand, when males were subjected to mild heat stress of the testis (43°C for 25 min), germ cells with inactivated *Mgat4d* were markedly more sensitive to the effects of heat stress, and transgenic mice expressing *Mgat4d* were partially protected from heat stress. Germ cells lacking *Mgat4d* generally mounted a similar heat shock response to control germ cells, but could not maintain that response. Several pathways activated by heat stress in wild type were induced to a lesser extent in *Mgat4d*[−/−] heat-stressed germ cells (NFκB response, TNF and TGFβ signaling, *Hif1α* and *Myc* genes). Thus, the Golgi glycoprotein MGAT4D is a novel, intrinsic protector of male germ cells from heat stress.

## Introduction

MGAT4D is designated family member D of the *MGAT4* gene family by the Human Genome Nomenclature Committee based on sequence similarity to other members, including MGAT4A and MGAT4B. The latter are N-acetylglucosaminyltransferases (GlcNAcTs) that add a β1,4GlcNAc to complex N-glycans. However, when MGAT4D is transfected into cultured cells, it does not appear to have GlcNAcT activity. Rather, it inhibits MGAT1 activity, the GlcNAcT responsible for initiating complex N-glycan synthesis ^1^. Because of this inhibitory activity, the protein was termed GnT1IP for GlcNAcT1 Inhibitory Protein. The *Mgat4d* gene is highly expressed in mouse testis with little expression in other mouse tissues ^2^. Based on RNA-seq analysis, it is expressed in spermatocytes and spermatids, but not in spermatogonia, sperm or Sertoli cells ^3^. MGAT4D is the most abundant protein in purified Golgi from rat testis germ cells ^4^. Characterization of the interactions of MGAT4D in the Golgi using a fluorescence resonance energy transfer (FRET) assay showed that it interacts with MGAT1 but not MGAT2, MGAT3, MGAT4B or MGAT5 ^3^. Since knockout of *Mgat1* in spermatogonia disrupts spermatogenesis and results in infertility ^5,6^, deletion or overexpression of *Mgat4d* in germ cells were both expected to have effects on spermatogenesis. In this paper, we show that unexpectedly, deletion of *Mgat4d* globally, or specifically in spermatogonia, or mis-expression of *Mgat4d* in spermatogonia, spermatocytes or spermatids, do not appear to alter spermatogenesis in young or aged mice, and do not affect fertility. However, mild heat stress of the testis in aged mice revealed that germ cells lacking *Mgat4d* exhibited more damage and apoptosis following heat stress. By contrast, a *Mgat4d* transgene expressed in spermatogonia, spermatocytes or spermatids, conferred partial resistance to mild heat stress. This is the first report of a germ cell intrinsic molecule that protects germ cells from heat stress and a novel function for a Golgi glycoprotein. Gene expression analyses showed that germ cells lacking *Mgat4d* responded to heat stress by initially upregulating heat shock and related genes. However, in contrast to controls, germ cells lacking *Mgat4d* did not sustain this response, nor upregulate anti-inflammatory and anti-apoptotic protective genes to the same degree as wild type germ cells. The data identify a new function for MGAT4D as a protector of male germ cell homeostasis, and provide new insight into how male germ cells withstand heat stress.

## Results

### Effects of global and conditional deletion of *Mgat4d* on spermatogenesis and fertility

Embryonic stem cells (ES Cells) carrying the construct *Mgat4d*^tm1a(KOMP)Wtsi^ designed to conditionally delete exon 4 of the *Mgat4d* gene (Fig. 1) were obtained from the Knockout Mouse Project (KOMP) repository. Following injection into C57BL/6J blastocysts, chimeras were crossed to C57BL/6J to obtain mice carrying the conditional *Mgat4d*^tm1a(KOMP)Wtsi^ allele. Male progeny were crossed with FVB *Stra8*-iCre ^7^ or *Flp1*-Cre transgenic females (129S4/SvJaeSor-*Gt(ROSA)26Sor*^*tm1(FLP1)Dym*^/J) ^8^. *Stra8* is expressed in spermatogonia from 3 days post-partum (dpp) and the *Flp1*-Cre was expressed from the ROSA26 locus. Male mice with global (*Mgat4d*[−/−]) or conditional (*Mgat4d*[F/F]:Stra8-iCre) inactivation of the *Mgat4d* gene were generated, and males expressing *LacZ* from the *Mgat4d* promoter were also obtained (Fig. 1). Both strains were crossed to FVB mice and maintained on a FVB background because *Mgat1* deletion was performed on the FVB background ^5^. Genotyping PCR identified *Mgat4d*[+], *Mgat4d*[−], *Mgat4d*[F] alleles and Stra8-iCre (Fig. 1). Primer sequences, locations and expected product sizes are given in Supplementary Table S1. Polyclonal rabbit antibodies (pAb) prepared against a C-terminal peptide of MGAT4D identified the long form (MGAT4D-L) and the short form (MGAT4D-S) which lacks 44 amino acids at the N-terminus of MGAT4D-L, and mice with inactivated *Mgat4d* had no signal, as expected (Fig. 1). Detection of LacZ expression by beta-galactosidase activity showed that the *Mgat4d* promoter is active mostly in spermatocytes and spermatids in testis tubules (Fig. 1), consistent with results of RNA-seq analysis ^3^. Immunohistochemistry for MGAT4D on testis sections from *Mgat4d*[+/−] or wild type males shows staining in the Golgi of spermatocytes and round spermatids, but not in spermatogonia or spermatozoa (Fig. 1), as observed in rat testis ^4^. Testis sections from *Mgat4d*[−/−] males showed no staining, as expected (Fig. 1).

**Figure 1.**
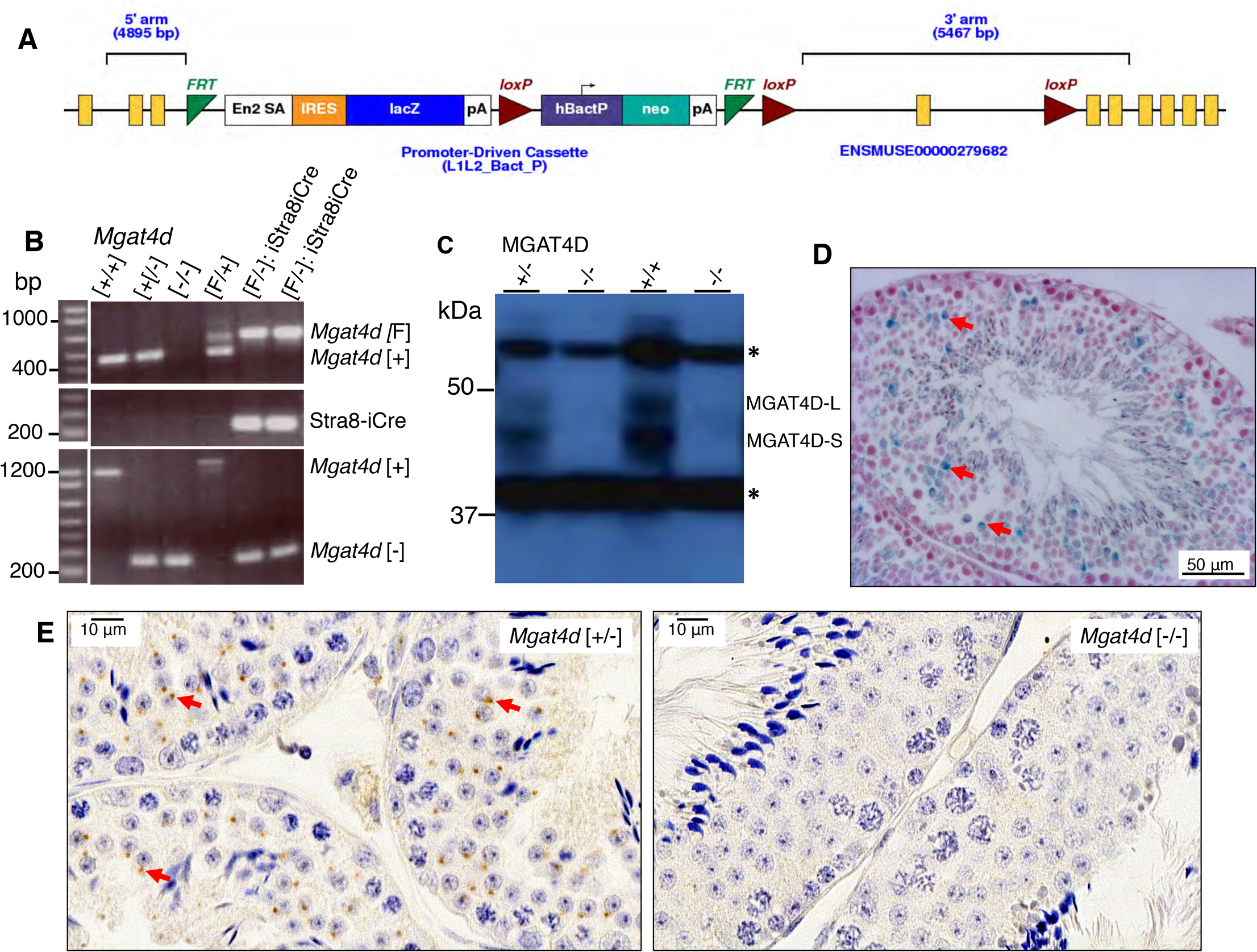
Generation of *Mgat4d* mutant mice. (**A**) Map of the targeted *Mgat4d*^tm1a(KOMP)Wtsi^ allele in ES cells obtained from KOMP. Exon 4 is flanked by two *loxP* sites. LacZ and the neomycin cassettes are flanked by two *Frt* sites. (**B**) PCR of genomic DNA from *Mgat4d*[+/+], *Mgat4d*[+/−], *Mgat4d*[−/−] and *Mgat4d*[F/−]:Stra8-iCre pups to determine genotype. Primers are given in Supplementary Table S1. (**C**) Western blot analysis of protein extracts of germ cells purified from 28 dpp *Mgat4d*[+/+], *Mgat4d*[+/−] and *Mgat4d*[−/−] mice. Long and short forms of MGAT4D are identified. * is a non-specific band (**D**) Representative testis section from a mouse carrying the *LacZ* gene under the control of the *Mgat4d* promoter after staining for β-galactosidase (blue). Nuclei were stained with eosin. (**E**) Immunohistochemistry of representative testis sections from *Mgat4d*[+/−] and *Mgat4d*[−/−] mice of 28 dpp. The presence of MGAT4D is shown by the brown stain consistent with a Golgi localization (arrows). Nuclei were stained with hematoxylin.

*Mgat4d*[−/−] males and females were fertile and transmitted the inactivated gene according to the expected Mendelian distribution (Table 1). Male mice with conditional deletion of *Mgat4d* in spermatogonia also showed no defects in fertility on a FVB background, or after backcrossing 10 generations to C57BL/6J mice (Table 1). Based on histological analyses, testicular weight and analysis of sperm parameters (sperm count, viability, morphology, motility and acrosome reaction), no obvious defects in spermatogenesis were observed in *Mgat4d*[−/−] males. In addition, aging (up to 596 dpp for FVB and 482 dpp for C57BL/6J) did not reveal apparent histological differences in spermatogenesis between mutant and control males (data not shown).

**Table 1.** Fertility of *Mgat4d*[−/−] and *Mgat4d-L-Myc* transgenic male mice

| Mouse strain | Male genotype | No. males | No. litters | No. pups | *Mgat4d*<sup>-/-</sup> | *Mgat4d*<sup>+/-</sup> | Transmission (CHI squared) |
| --- | --- | --- | --- | --- | --- | --- | --- |
| FVB | *Mgat4d*<sup>-/-</sup> | 5 | 5 | 54 | 29 | 25 | 0.586 |
| C57BL/6J | *Mgat4d*<sup>-/-</sup> | 4 | 13 | 88 | 42 | 46 | 0.669 |
| C57BL/6J | *Mgat4d*<sup>[F/F]:Stra8-iCre</sup> | 2 | 9 | 59 | 29 | 30 | 0.896 |
|  |  |  |  |  | *Mgat4d-L-Myc* | *Mgat4d*<sup>[+/+]</sup> |  |
| C57BL/6J | *Stra8-Mgat4d-L-Myc* | 7 | 20 | 171 | 70 | 101 | 0.018 |
| C57BL/6J | *Ldhc-Mgat4d-L-Myc* | 4 | 9 | 59 | 26 | 33 | 0.362 |
| C57BL/6J | *Prm1-Mgat4d-L-Myc* | 6 | 12 | 77 | 41 | 36 | 0.568 |

*Mgat4d*<sup>-/-</sup> males were crossed with *Mgat4d*<sup>+/-</sup> females. *Mgat4d*<sup>[F/F]:Stra8-iCre</sup> males were crossed with *Mgat4d*<sup>[F/-]</sup> females. *Mgat4d-L-Myc* transgenic heterozygote males were crossed with *Mgat4d*<sup>[+/+]</sup> females.

As discussed in the Introduction, MGAT4D was initially described as an inhibitor of MGAT1 activity and termed GnT1IP ^1^. By deleting such an inhibitor, we expected MGAT1 activity might increase, and the level of complex N-glycans on glycoproteins might also increase. We determined MGAT1 GlcNAc transferase activity in germ cell extracts. Germ cells were purified from 28 dpp C57BL/6J wild type (n=4) and *Mgat4d*[−/−] males (n=4) and protein extracts prepared. The average activity for *Mgat4d*[+/+](1.86+/−0.38 nmol/mg/hr) and for *Mgat4d*[−/−](1.68+/−0.32 nmol/mg/hr) were not significantly different (*p*=0.72). This result might reflect the fact that *Mgat1* is most highly expressed in spermatogonia which do not express *Mgat4d* ^3^. However, there was no evidence of a specific increase in complex N-glycan species in *Mgat4d*[−/−] testis sections subjected to MALDI mass spectrometry imaging (MALDI-IMS) for N-glycans (Supplementary Fig. S1).

### Males lacking *Mgat4d* are more sensitive to mild heat stress of the testis

Given the apparent lack of significant consequences for spermatogenesis of removing *Mgat4d*, we investigated whether stressing testicular germ cells would reveal any effects of *Mgat4d* loss. Spermatogenesis is sensitive to an increase in temperature ^9,10^ and we reasoned that disturbing tissue homeostasis using mild heat stress might reveal roles for MGAT4D in testis. The remaining cohort of aged *Mgat4d*[+/−] and *Mgat4d*[−/−] FVB mice of between 592 and 596 dpp were anesthetized and subjected to mild heat stress by immersing the lower half of the body in water at 43°C for 25 min. Mock treatment involved the same procedure with a water temperature of 33°C. After recovery for 24 hr, testes were harvested. One testis was used for histological analysis and the other for RNA and protein extraction. While testis sections from males treated at 33°C appeared normal, 43°C treatment caused the appearance of enlarged (≥10 μm) multinucleated cells, large vacuoles (≥10 μm), small vacuoles and pyknotic cells in testis tubules (Fig. 2). Spermatozoa in the epididymis also included pyknotic cells following heat stress (Fig. 2). Compared to controls, *Mgat4d*[−/−] testis sections exhibited an increased number of tubules (~3.5-fold) with enlarged cells, and a decrease in undamaged tubules (~2-fold). (Fig. 2). No significant difference was found in testis weights of heat-treated versus control mice (Supplementary Table S2).

**Figure 2.**
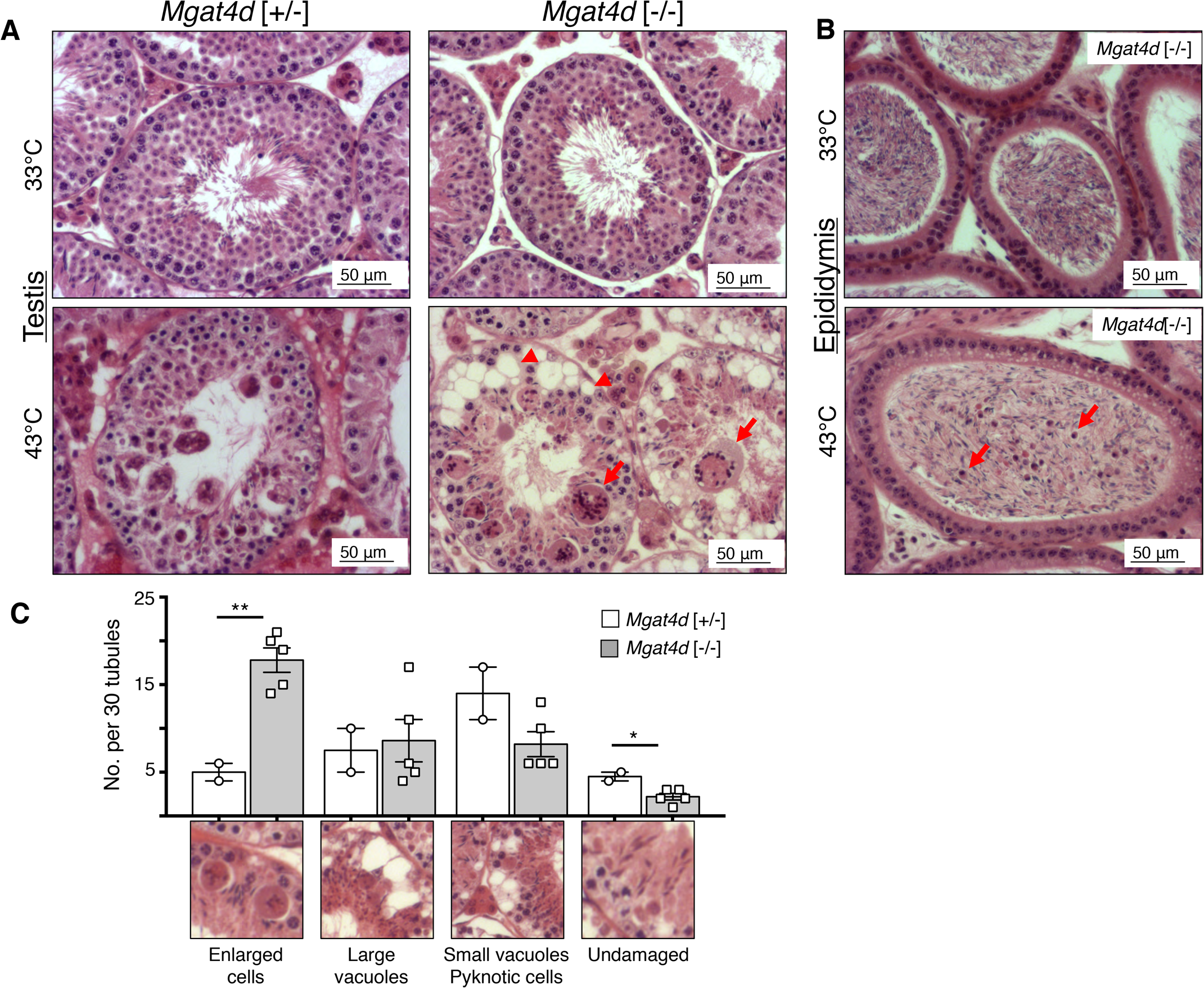
Effects of heat treatment on *Mgat4d*[−/−] testes. (**A**) Representative testis sections stained with H&E. Upper panels from mice whose lower body was submerged for 25 min at 33°C and lower panels from mice treated similarly at 43°C. Arrows indicate enlarged cells, arrow heads show vacuoles in germ cells. (**B**) Representative epididymis sections from a *Mgat4d*[−/−] male treated at 33°C (upper) or 43°C (lower) and stained with H&E. Arrows in the 43°C sample indicate pyknotic cells in the tubule lumen. (**C**) Quantification of different tubule categories in testis sections from heat-treated (43°C) *Mgat4d*[+/−] and *Mgat4d*[−/−] males. Positive tubules were counted as those with at least one cell of radius ≥10 μm; large vacuoles were tubules with at least one vacuole ≥10 μm; small vacuoles, pyknotic cells were tubules with at least one vacuole of radius <10 μm or tubules containing pyknotic cells; undamaged tubules were tubules with no apparent damage. Mice were from an aged cohort (592-596 days) of *Mgat4d*[+/−](n=2) and *Mgat4d*[−/−](n=5) mice. Thirty (30) tubules were counted in one section per mouse. Student’s t test (two-tailed, unpaired) **p<0.01; *p<0.05.

Heat stress increases apoptosis in differentiating germ cells ^10–12^ and so testis sections from heat- and mock-treated aged FVB males were subjected to the “Apoptag” assay and staining was quantified using FIJI software (https://fiji.sc/). As expected, apoptosis increased in sections from control heat-treated males sacrificed 24 hr after heat treatment (Fig. 3). However, testes from *Mgat4d*[−/−] mice showed ~2-fold more apoptotic germ cells than *Mgat4d*[+/−] controls (Fig. 3). Thus, based on histology and levels of apoptosis, the effects of heat stress were more severe for aged *Mgat4d*[−/−] testes than for heterozygous testes.

**Figure 3.**
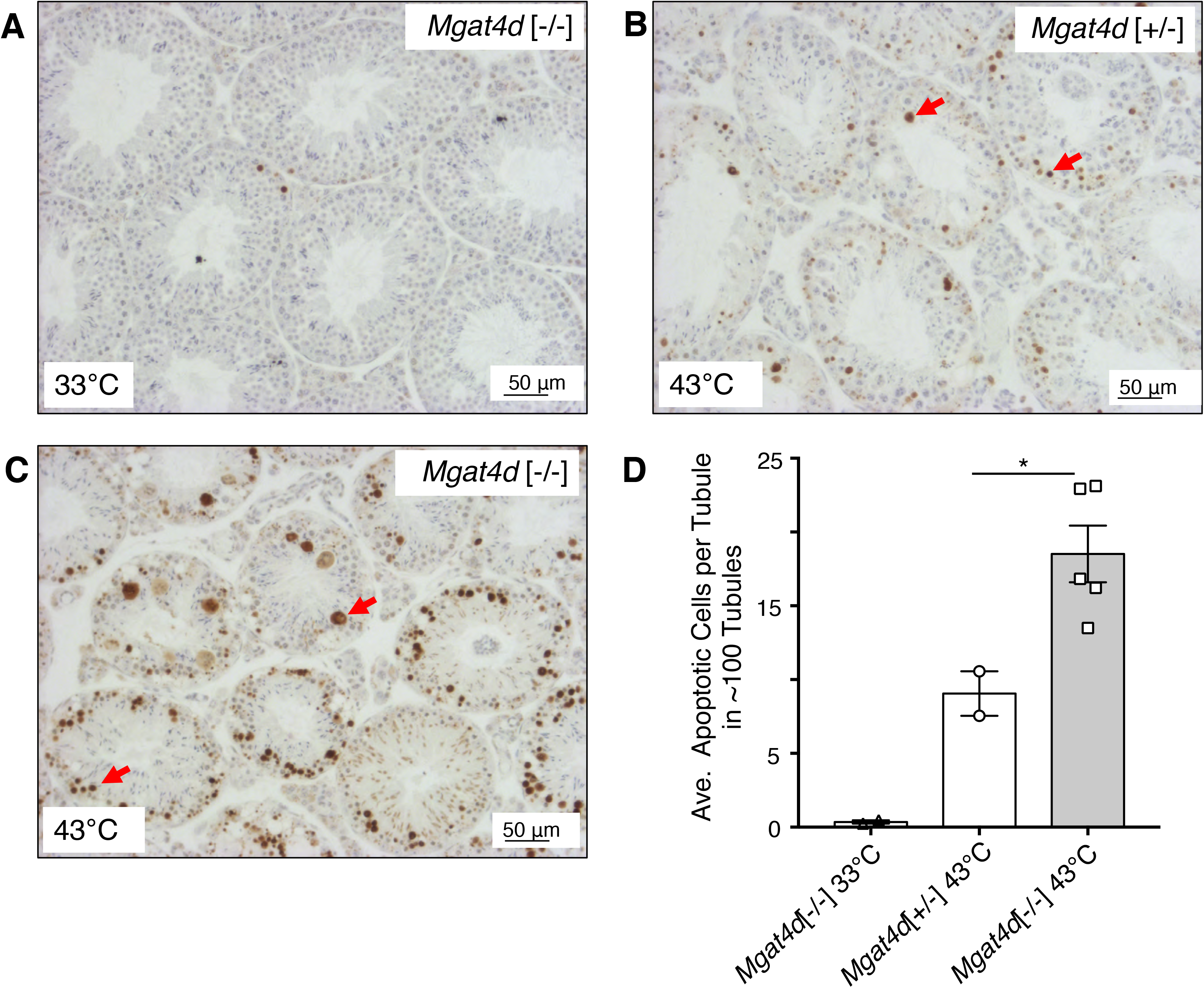
Apoptosis of germ cells in heat-treated testes. Representative testis sections from 33°C- or 43°C-treated aged FVB males were subjected to the TUNEL “Apoptag” assay for in situ detection of DNA strand breaks. (**A**) Section from a 33°C-treated *Mgat4d*[−/−] male. (**B**) Section from a 43°C-treated *Mgat4d*[+/−] male. (**C**) Section from a 43°C-treated *Mgat4d*[−/−] male. DNA breaks stained brown (red arrows). (**D**) quantification of apoptotic signal in ≥100 tubules using FIJI software. Student’s t test (unpaired, two-tailed) *p<0.05.

### *Mgat4d* transgenic mice are resistant to the effects of heat stress

Mice with targeted deletion of *Mgat1* in testicular germ cells exhibit defective spermatogenesis and are infertile ^5^. Thus, it was expected that inhibiting MGAT1 activity by increasing the level of MGAT4D in germ cells, would induce defects in mouse spermatogenesis. To investigate, C57BL/6J transgenic males expressing a *Mgat4d-L-Myc* cDNA in specific germ cell types were generated. This transgene has previously been shown to inhibit MGAT1 in transfected cells ^1,3^. The *Stra8* (Stimulated By Retinoic Acid 8) promoter was used to express the transgene in spermatogonia ^5–7^, the *Ldhc* (Lactate Dehydrogenase C) promoter was used to express in spermatocytes ^13,14^, and the *Prm1* (Protamine 1) promoter was used to express in spermatids ^15^ (Fig. 4). The transgenic mouse strains were named *Stra8-Mgat4d-L-Myc*, *Ldhc-Mgat4d-L-Myc* and *Prm1-Mgat4d-L-Myc*, respectively. They were genotyped by PCR of genomic DNA using primers described in Supplementary Table S1, and transgene expression was shown to be 3-6-fold greater than endogenous *Mgat4d-L* levels using quantitative RT-PCR (qRT-PCR) on cDNA from testis (Fig. 4). qRT-PCR using primers specific for the *Myc* sequence gave a similar level of expression based on Ct values (not shown). Myc transcripts could not be quantitated relative to the control that has no transgene. By contrast, attempts to determine MGAT4D-L-Myc protein levels in testis extracts by western blot analysis using anti-Myc monoclonal antibodies (mAb) from several species were not successful, although MGAT4D-L-Myc overexpressed in CHO cells is detected by anti-Myc mAb ^3^. We generated C- and N-terminal peptide-purified rabbit pAbs that detect MGAT4D-L-Myc or Myc-MGAT4D-L, respectively, in transfected CHO cells (Supplementary Fig. S2). The C-terminal pAb detected Myc-MGAT4D-L much more readily than MGAT4D-L-Myc (Supplementary Fig. S2). Moreover, MGAT4D-L-Myc was not detected in extracts from transgenic germ cells. Nevertheless, the results that follow show that each transgene was functional in the heat stress test.

**Figure 4.**
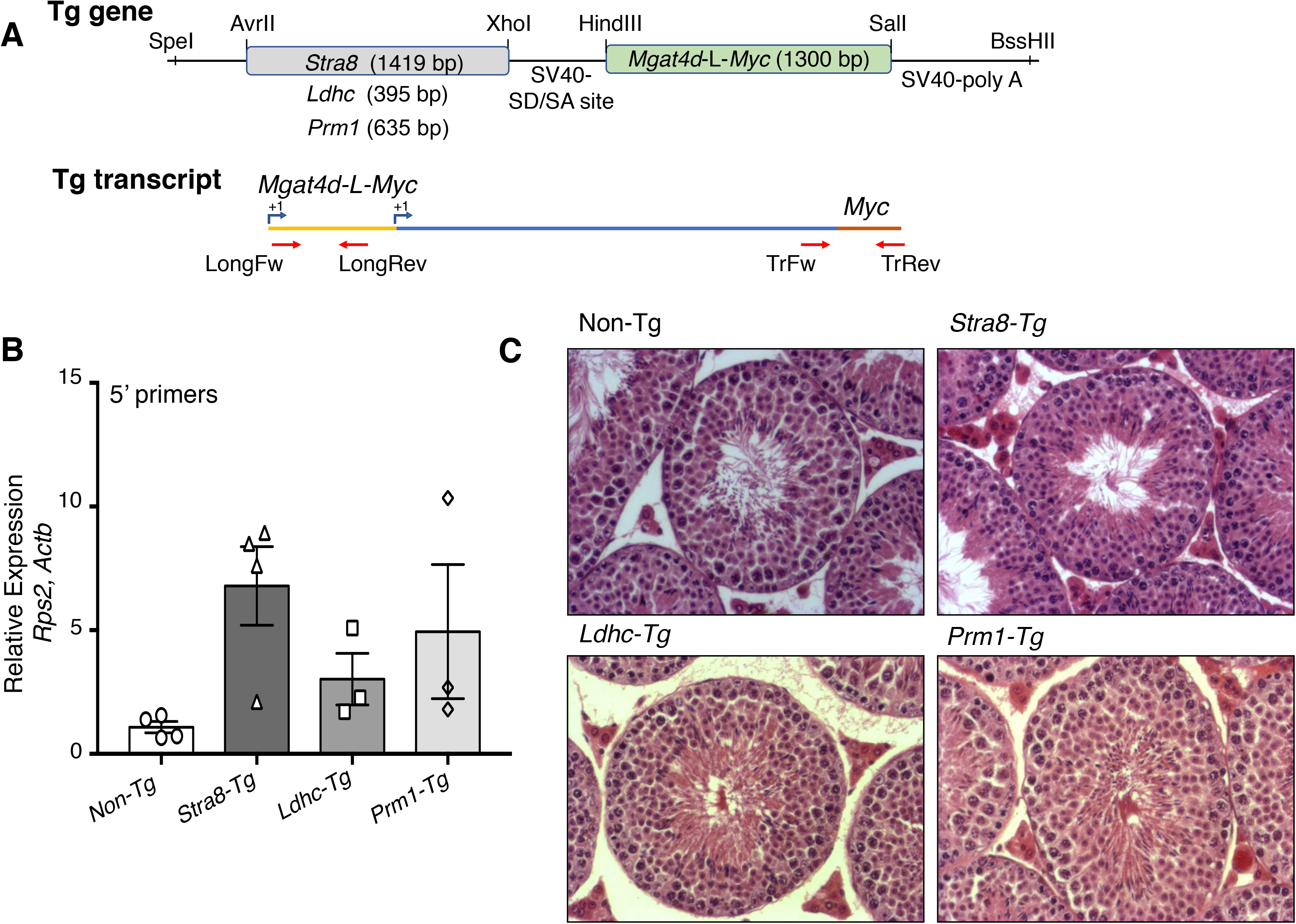
Generation and characterization of *Mgat4d* transgenic mice. (**A**) Schematic representation of constructs used to generate transgenic (Tg) mice. Expression of *Mgat4d-L-Myc* was driven by promoters (*Stra8, Ldhc* and *Prm1*) specific for different germ cell types. Lower diagram, position of primers used for qRT-PCR amplification. “LongFw” and “LongRev” to amplify the 5’ region of Mgat4d-L; “TrFw” and “TrRev” to amplify the transgene junction. (**B**) qRT-PCR of *Mgat4d*-L 5’ primer transcripts relative to *Actb* and *Rps2*. Testis RNA was isolated from males of 28 dpp. (**C**) Representative H&E stained testis sections from 120 dpp control and *Mgat4d-L-Myc* transgenic males.

Overexpression or mis-expression of *Mgat4d* in germ cells was expected to inhibit MGAT1 activity ^1,3^. However, compared to *Mgat4d*[+/+] controls (1.1 +/−0.13 nmol/mg/hr; n=10), there was no significant inhibition of MGAT1 activity in germ cells from 28-42 dpp *Stra8-Mgat4d-L-Myc* (1.4+/−0.16 nmol/mg/hr; n=7)*, Ldhc*-*Mgat4d-L-Myc* (0.78 +/−0.18 nmol/mg/hr; n=4)*, Prm1-L*-*Mgat4d* (1.3 +/−0.03 nmol/mg/hr; n=4). In addition, the MALDI-IMS of testis sections showed that the complement of complex N-glycan species was not reduced in testis sections from transgenic mice (Supplementary Fig. S1). Rather, there was a significant increase in both oligomannosyl and simple complex N-glycans.

Histological analysis of testis sections showed no obvious changes in spermatogenesis or testicular structure in adult transgenic mice (Fig. 4). In addition, the fertility of transgenic males was normal, although *Stra8-Mgat4d-L-Myc* mice showed low transgene transmission from transgenic males (Table 1). Males from the three transgenic mouse strains and non-transgenic littermates or wild type C57BL/6J controls were subjected to mild heat stress. No significant difference was observed in testis weights of mock-versus heat-treated mice (Supplementary Table S2). Importantly, however, each transgenic strain showed an ~3-fold reduction in the number of tubules with enlarged germ cells, and ~2-fold fewer had tubules with large vacuoles (Fig. 5). The number of undamaged tubules was also increased but small vacuoles and pyknotic cells were present in heat-treated transgenic mice (Fig. 5). The “Apoptag” assay revealed an ~2-fold reduction in apoptotic germ cells in all three transgenic strains (Fig. 6). We also investigated previously reported gene expression changes due to heat stress. In wild type males, *Socs3, Hspa1a* and *Degs1* were up-regulated, while *Bcl2l12, Crbg3* and *Dmrt1* were down-regulated after heat treatment (Fig. 7), consistent with previous observations ^11,16^. Interestingly, *Mgat4d* was markedly down-regulated following heat treatment (Fig. 7). Gene expression in heat treated *Stra8-Mgat4d-Myc* males was similar to that in non-transgenic males at 33°C, up-regulated genes being less up-regulated and down-regulated genes, less down-regulated compared to non-transgenic at 43°C (Fig. 7). Thus, on the basis of several criteria, the presence of a *Mgat4d-L-Myc* transgene in germ cells gave significant protection from heat stress.

**Figure 5.**
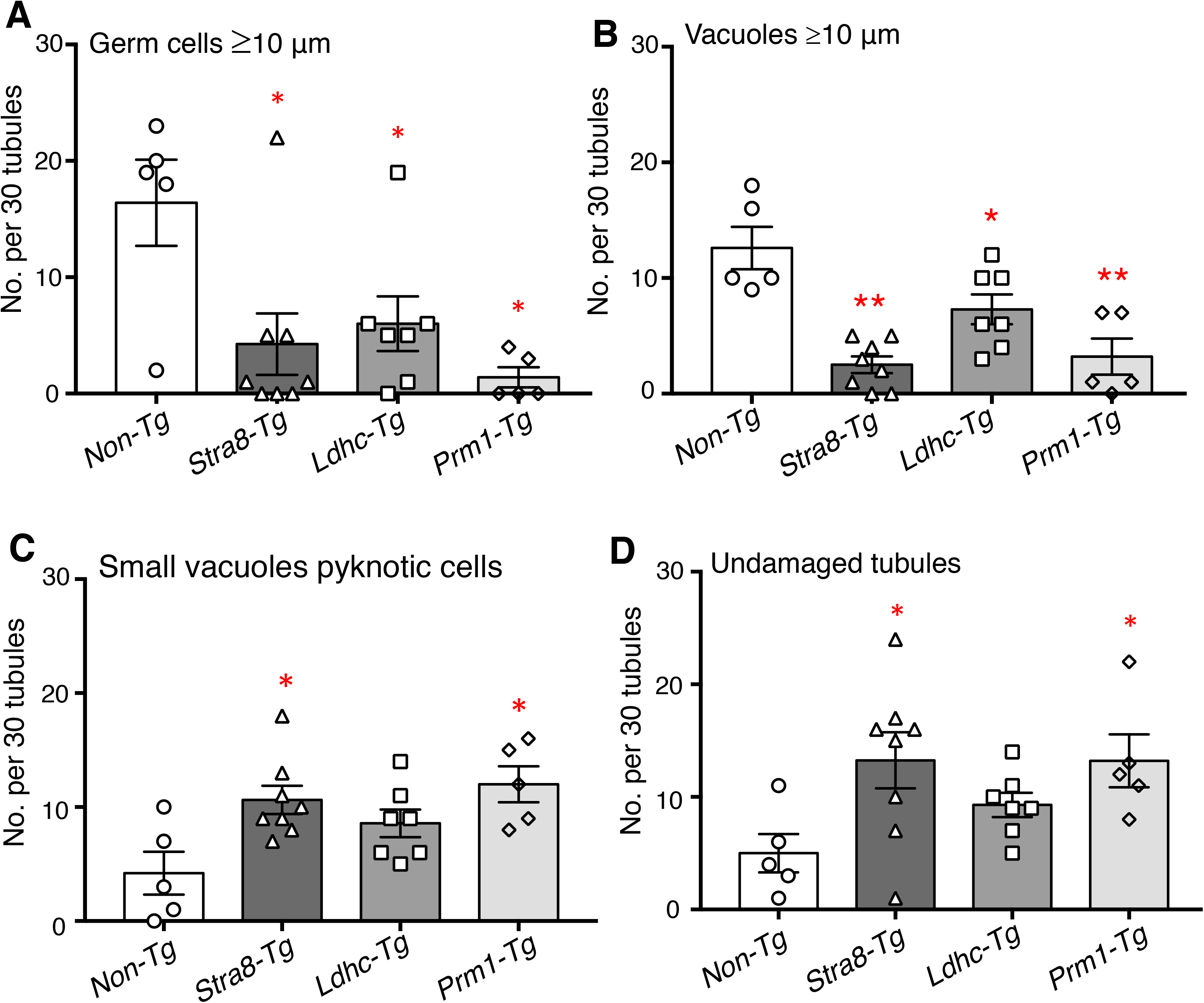
Effects of heat stress in *Mgat4d* transgenic testis. Quantification of heat-induced damage in testes of *Mgat4d*[+/+] (n=5), *Stra8-Mgat4d-L-Myc* (n=8), *Ldhc-Mgat4d-L-Myc* (n=7), and *Prm1-Mgat4d-L-Myc* (n=5) C57BL6/J males. 30 tubules were investigated per mouse in one H&E stained testis section to detect (**A**) Enlarged germ cells; (**B**) Large vacuoles; (**C**) Small vacuoles, pyknotic cells; (**D**) Undamaged tubules. Differences from control two- tailed, unpaired Student t-test *p<0.05, **p<0.01.

**Figure 6.**
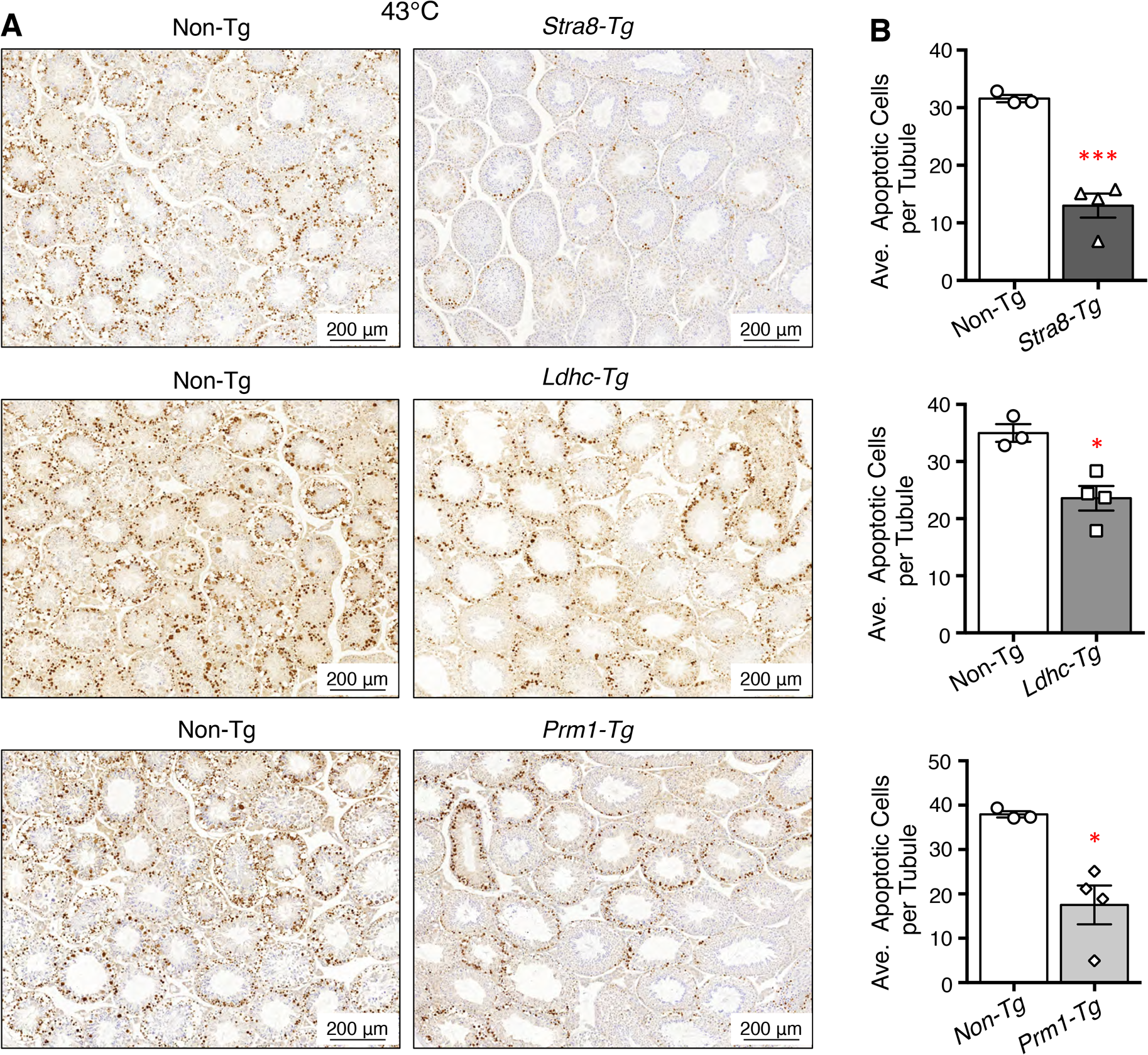
Effects of heat treatment on apoptosis in *Mgat4d* transgenic mice. (**A**) Representative images of testis sections from 43°C-treated *Mgat4d* [+/+] (n=3) and *Mgat4d* transgenic mice (n=4 for each) stained by the Apoptag kit to detect DNA breaks. (**B**) Quantification of Apoptag signal in ≥100 tubules of 43°C-treated non-transgenic and transgenic mice. Statistical differences determined by two-tailed unpaired Student’s t-test *p<0.05, ***p<0.001.

**Figure 7.**
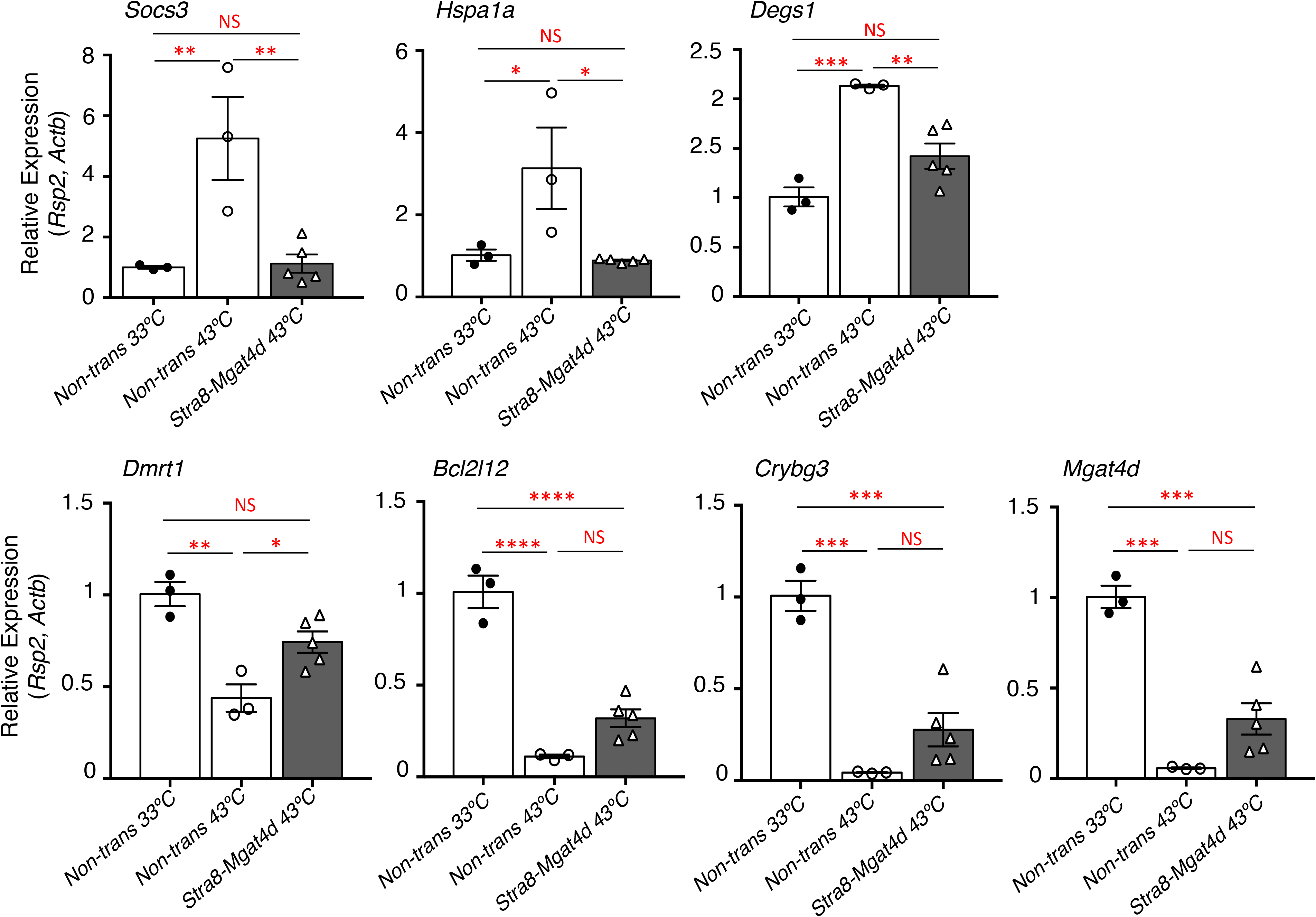
qRT-PCR of cDNA from testis of 33°C- or 43°C-treated control vs *Stra8-Mgat4d-L-Myc* males of 7 months. Testes were isolated 24 hr after treatment. qRT-PCR was performed in triplicate. Relative gene expression was normalized to *Actb* and *Rps2*. Mean ± SEM; statistical analysis by two-tailed, unpaired Student’s t-test.; *p<0.05, **p<0.01, ***p<0.001, ****p<0.0001.

### Molecular basis of the increased sensitivity of *Mgat4d*[−/−] germ cells to heat stress

The histological and apoptotic changes induced by heat stress reported above were observed in an aged cohort of 1.6 year FVB mice. We subsequently tested 7 month C57BL/6J *Mgat4d*[−/−] mice and did not observe increased sensitivity to heat stress. However, protection from heat stress was observed in adult C57BL/6J transgenic mice as shown here. Thus, to determine whether *Mgat4d*[−/−] mice on a C57BL/6J background exhibited a more sensitive response to heat stress than controls, and to also gain insights into molecular mechanisms that underlie this phenotype, microarray analyses were performed on cDNA from purified germ cells of control and *Mgat4d[−/−]* C57BL/6J mice of 2 months. *Mgat4d*[+/+] and *Mgat4d*[−/−] males were treated at 33°C or 43°C for 25 min and sacrificed after 8 hr, a time when no visible histological changes to germ cells were observed (data not shown). Testes were enzymatically dissociated and germ cells were isolated and counted. RNA preparations with a RIN value >7.9 were used to make cDNA for microarray analysis. Purity of germ cells was assessed by relative expression of germ cell-specific and non-germ cell genes to the same genes expressed in testis RNA as previously described ^6^. The Mouse Clariom™ D GeneChip™ Mouse Transcriptome Array 1.0 from Affymetrix was used. Custom scripts using the R/Bioconductor tools affymetrix and limma were used to process the raw (.CEL) files and to compare *Mgat4d*[−/−] versus *Mgat4d*[+/+] microarray data from 33°C- and 43°C-treated mice. The samples displayed a moderate clustering by genotype, as seen in PCA plots (Fig. 8). Importantly, significant differences between genotypes were much less pronounced at 33°C than at 43°C, as witnessed by the tighter correlation in the heat maps, and a lower number of differentially expressed genes (DEGs) between genotypes shown in volcano plots (Fig. 8). However, a clear difference was evident between wild type and *Mgat4d*[−/−] arrays from germ cells of mice treated at 43°C. Given the importance of the temperature as a confounding variable, it was included in modelling differential gene expression between genotypes. DEGs in mutant versus wild type germ cells at 33°C and at 43°C were determined, and the interaction between temperature and genotype was evaluated to obtain gene lists for further analysis. Microarray data are deposited in NCBI’s Gene Expression Omnibus (GEO) and are accessible through GEO serial accession number GSE137307.

**Figure 8.**
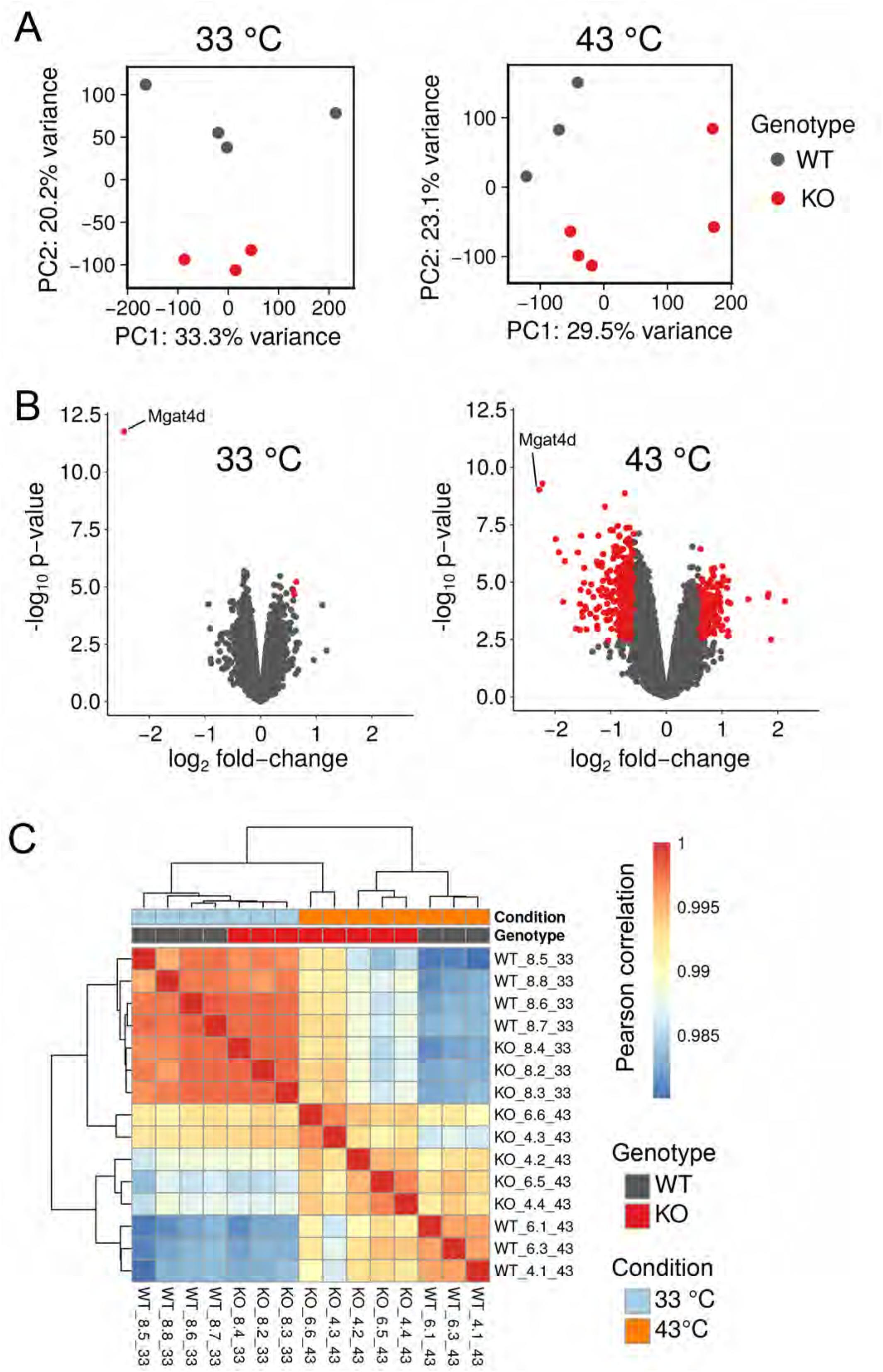
Microarray analysis of germ cell cDNA. Two month C57BL/6J males were treated at 33°C or 43°C for 25 min and germ cells were harvested 8 hr after recovery. (**A**) Principal component analysis (PCA) for 33°C-treated *Mgat4d*[+/+] (n=4) and *Mgat4d*[−/−] (n=3) arrays and 43°C-treated *Mgat4d*[+/+] (n=3) and *Mgat4d*[−/−] (n=5) samples. (*B*) Volcano plots showing the distribution of DEGs based on their expressed log_2_ fold-change and −log_10_ *p* value. Red dots represent genes with a log_2_ fold-change lower than −0.6 or higher than 0.6 (equivalent to +/− 1.5 fold-change) and a *p* value below the threshold of 0.05 (−log10 (0.05) = 1.3). (**C**) Correlation analysis of microarray chip data of wild type (WT) and *Mgat4d*[−/−] (KO) germ cells at 33°C and 43°C.

Analysis of 33°C *Mgat4d*[−/−] versus control microarrays with FDR<0.05 and fold change +/−1.5 gave 4 DEGs (3 up- and 1 down-regulated gene), including *Mgat4d* as expected. *Mgat4d* transcripts were not completely lacking in *Mgat4d*[−/−] samples due to transcription beyond the deleted exon 4 (Supplementary Fig. S3). However, no MGAT4D protein was detected by western analysis or immunohistochemistry (Figure 1, Supplementary Fig. S2). One of the up-regulated genes, pseudogene Gm12584, maps to the locus of a testis-specific gene, adenosine deaminase domain-containing 1 (*Adad1*), which encodes a nuclear RNA-binding protein ^17^. Upregulated Gm24265 refers to an SnRNA mapped to chromosome 5. The third up-regulated gene was *Tmed6* (Transmembrane P24 Trafficking Protein 6) which is enriched in the endoplasmic reticulum and Golgi compartments. It is notable that all the upregulated DEGs have a log fold change <1 (fold change <2), revealing a very mild effect of *Mgat4d* deletion on germ cell gene expression under control conditions (Supplementary Table S3). By contrast, analysis using the same stringency for data from heat-stressed *Mgat4d*[−/−] versus control germ cells, revealed 476 DEGs in *Mgat4d*[−/−] germ cells with 110 genes up-regulated and 366 genes down-regulated. The top down-regulated genes were *Serpinb1a* (2.13 log fold change) followed by *Ly96* (2.12 Log fold change) and S100-a11 (2.09 Log fold change) (Supplementary Table S4). Some of the down-regulated genes (*Serpinb1a, Star, Osr2, Klk1b22, Itih2*) are related to the regulation of cellular homeostasis, proliferation or survival ^18–23^. The top two up-regulated genes were non-coding Gm26715 and Gm48565 (2.04 and 1.98 log fold change respectively), followed by *Hspa1a* and *Hspa1b* heat shock proteins (1.81 and 1.75 log fold change, respectively). Most of the up-regulated genes were non-coding or predicted genes (Supplementary Table S5). cDNA from the 43°C-treated control and *Mgat4d*[−/−] cDNA preparations was used for qRT-PCR validation of DEGs observed in microarray experiments (Fig. 9). The relevant primer sequences are given in Supplementary Table S6.

**Figure 9.**
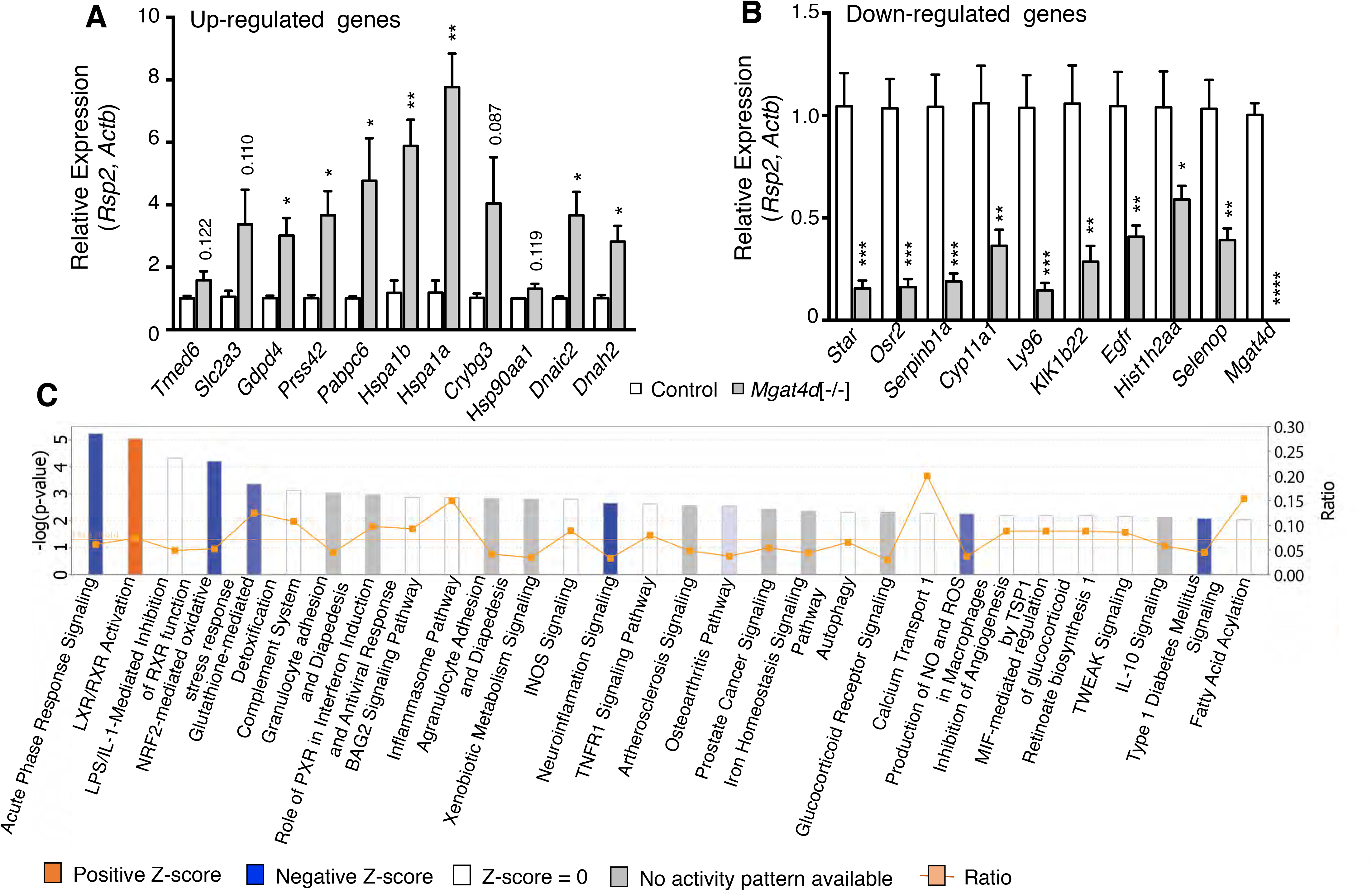
Validation and Ingenuity Pathway analysis (IPA). (**A**) qRT-PCR validation of up-regulated genes in control (n=3) and *Mgat4d*[−/−] (n=4) cDNA samples from the same mice used for microarray analyses. (**B**) qRT-PCR of down-regulated genes in the same samples. Relative expression was determined using *Actb* and *Rps2*. Assays were performed in triplicate. Error bars represent mean ± SEM; statistical analysis by two-tailed, unpaired Student’s t-test \**p*<0.05, \*\**p*<0.01, \*\*\**p*<0.001. (**C**) Top canonical pathways in IPA significantly overrepresented in heat-treated *Mgat4d*[−/−] germ cells compared to wild type, normalized to their respective 33°C counterparts, according to −log *p* value.

### Enriched biological pathways in heat-stressed *Mgat4d*[−/−] germ cells based on Ingenuity Pathway Analysis (IPA)

To find the most significantly represented pathways differentially altered in *Mgat4d*[−/−] versus *Mgat4d*[+/+] germ cells following heat stress, we examined the relationship between DEGs at a ±1.5 fold change with adjusted FDR<0.05 and *p*<0.05 using IPA. Interestingly, the top canonical pathways were mostly down-regulated or with “no activity pattern available” in 43°C-treated *Mgat4d*[−/−] versus control germ cells, and were related to recovery from stress conditions (Fig. 9). Ranked by −log(p-value), the top down-regulated pathway was Acute Phase Response Signaling (*p*=5.2; Z-score −2.45) followed by LXR/RXR Activation (*p*=5.04; Z-score 0.707), the only pathway with a positive z-score and −log(p-value) higher than 2. NRF2-mediated Oxidative Stress Response (*p*=4.98; Z-score −2.111) and Glutathione-mediated Detoxification (*p*=3.37; Z-score −2) were also top pathways.

### Top upstream transcriptional regulators

IPA was used to predict the top upstream transcriptional regulators in the DEGs based on their gene targets. The algorithm calculates a p-value on the basis of significant overlap between genes in our test dataset and target genes regulated by the same regulator in the IPA knowledge base. The activation Z score algorithm was used to make predictions. This analysis identified 323 upstream regulators with a p-value of overlap <0.05 and a Z-score greater than or equal to +/−2. *Tgfb1*, *Tnf*, *Ifng*, *Il1b* related to immune system regulation are the top inhibited upstream regulators (Supplementary Table S7). Sorting the results by Expression Log Ratio +/−1, identified 13 differentially-expressed upstream regulators in our data set, 11 down-regulated and 2 up-regulated (Supplementary Table S7).

### Most represented networks, toxicological functions, diseases and biological functions

DEGs in germ cells from heat-treated mice were compared by IPA with genes belonging to specific biological networks or implicated in diseases. The most highly ranked network was “DNA Replication, Recombination, and Repair, Nucleic Acid Metabolism, Small Molecule Biochemistry” with 28 focus molecules (Supplementary Table S8). The top diseases and biological functions were related to “Organismal Survival” - 19 biological functions were predicted to be increased with an activation Z-score between 6.131 and 2.01, mostly related to inflammation, injury and disease (Supplementary Table S9) but a higher number of diseases or functions were predicted to be decreased (71). The top category was “Lipid Metabolism, Small Molecule Biochemistry, Vitamin and Mineral Metabolism” and the most represented of these were related to cellular function.

### Gene set enrichment analysis (GSEA)

Comparisons of DEGs at 33°C and 43°C with published, classified gene sets in the MSigDB was performed using GSEA ^24–26^. Of the eight categories of gene sets, the Hallmark collection summarizes well-defined biological processes and states from v4.0 MSigDB collections C1 through C6 ^27^. Hallmark gene sets with a Normalized Enrichment Score (NES) of +/−2, FDR<0.25 and *p*<0.05 were examined. In *Mgat4d*[−/−] germ cells, only 3 Hallmark gene sets were significantly enriched at 43°C - E2F targets, G2M checkpoint and spermatogenesis. The Hallmark Spermatogenesis gene set contains genes upregulated during the process of spermatogenesis, indicating that loss of *Mgat4d* in heat-stressed germ cells leads to induction of spermatogenesis-promoting genes as a response, whereas germ cells expressing *Mgat4d* were comparatively protected from premature upregulation of these genes (Supplementary Fig. S4). Gene sets enriched in *Mgat4d*[+/+] germ cells at 43°C were related to immune pathways signaling, inflammatory responses, apoptosis and hypoxia (Supplementary Fig. S4).

Gene sets of note in other collections were: negative regulation of extrinsic apoptosis signaling (suppression of apoptosis) by *Mgat4d* in the C5 collection; increased inflammatory response and TNF targets up in *Mgat4d*[+/+] germ cells in the C2 collection; late ATM-dependent genes induced by radiation up in *Mgat4d*[+/+]; increased induction in *Mgat4d*[−/−] of MYBL1 target genes in spermatocytes; and genes downregulated in response to gamma-radiation were up in *Mgat4d*[−/−]. We also investigated DEGs in wild type versus mutant at 43°C versus 33°C using EnrichR ^28^. Heat maps highlight some of the informative EnrichR gene sets and also show illustrative gene expression differences identified (Fig. 10). The overall results suggest that *Mgat4d*[−/−] germ cells have a problem responding to heat shock stress, e.g. coping with hyperthermic stress through clearance of damaged proteins (*Casp8;* Fig. 10). A number of pathways and genes were induced to a lesser extent in *Mgat4d*[−/−] heat-stressed mice, including *Hif1α*, the NFκB response, pro-inflammatory pathways such as TNF and TGFβ signaling, and genes that promote proliferation such as *Myc* (Fig. 10).

**Figure 10.**
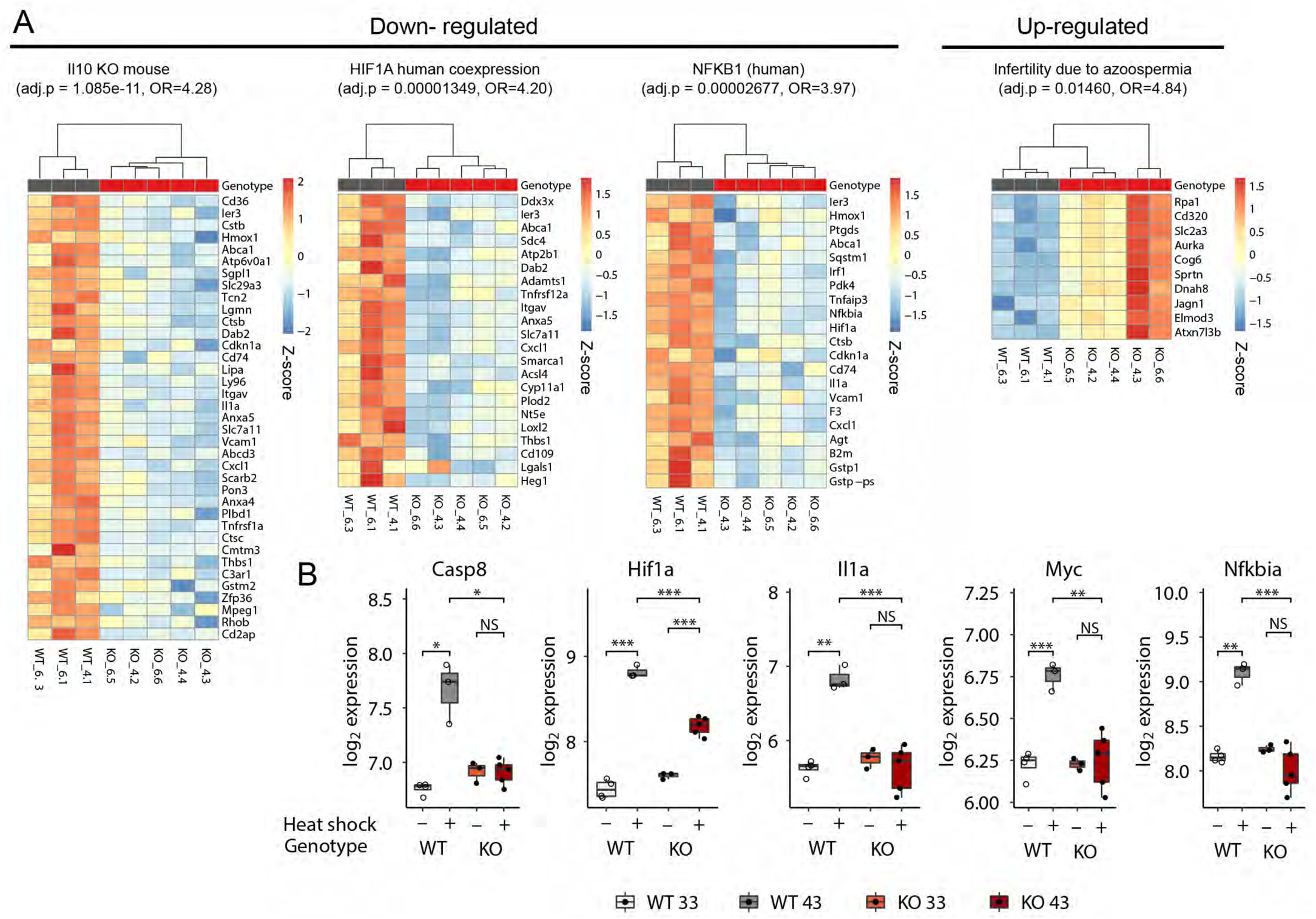
Differential gene interactions between *Mgat4d* genotype and heat shock conditions. (**A**) Heat maps showing DEGs either down- or up-regulated specifically in *Mgat4d* wild type, but not *Mgat4d* KO cells following heat shock, representative of significantly enriched pathways identified by EnrichR (http://amp.pharm.mssm.edu/Enrichr/). Adjusted *p* values and odds ratios (OR) for the respective pathways are shown. Full names of the pathways are: Single Gene Perturbations from GEO: Il10 KO mouse GSE25846 sample 3062; ARCHS4 TFs Coexp: HIF1A_human_tf_ARCHS4_coexpression; TRRUST Transcription Factors 2019: NFKB1_ human; Disease Perturbations from GEO up: Infertility due to azoospermia C1321542 mouse GSE3676 sample 151. Color scales represent gene-wise Z-scores. (**B**). Box plots showing expression of representative genes of the indicated pathways across genotypes and heat shock conditions. These genes were much more up-regulated by heat treatment in WT compared to mutant (KO) germ cells.

## Discussion

In this paper we characterize the first germ cell intrinsic molecule that protects from heat stress - the Golgi glycoprotein MGAT4D. Other molecules that protect germ cells from heat stress have been described, but each was overexpressed in testis under an exogenous promoter ^29,30^. MGAT4D maps to mouse chromosome 8 whereas previous genetic loci linked to germ cell resistance to heat stress map to mouse chromosomes 1 and 11 ^31^. Global deletion of the ion channel *Trpv1* increases the sensitivity of germ cells to heat stress ^32^, and this gene maps to chromosome 11, albeit 5.5cM away from the heat resistant locus on chromosome 11 ^31^. However, it is not clear which cells of the testis express *Trpv1* which is most highly expressed elsewhere in dorsal root ganglia. *Mgat4d* is most highly expressed in spermatocytes and spermatids ^3^ and thus well positioned to protect germ cells from heat stress. Here we provide several pieces of evidence in support of such a germ cell protective role for *Mgat4d*. First, we show that an old cohort of *Mgat4d*[−/−] males were more sensitive to mild testicular heat stress than heterozygote controls, as evidenced by increased germ cell defects and apoptosis at 24 hr after heat stress. Second, we found that mice expressing a *Mgat4d-L-Myc* transgene in either spermatogonia (*Stra8* promoter), spermatocytes (*Ldhc* promoter) or spermatids (*Prm1* promoter) were less sensitive to testicular heat stress than wild type controls, based on reduced germ cell defects and reduced apoptosis. Characterization of individual gene expression changes for genes known to exhibit increased or decreased expression following heat stress, showed that males expressing the *Stra8-Mgat4d-L-Myc* transgene were comparatively resistant to heat stress and, at 43°C, behaved similarly to non-transgenic germ cells treated at 33°C, whereas non-transgenic males treated at 43C showed the marked gene expression changes predicted from the literature. To investigate gene expression differences in more depth, we performed microarray analyses on *Mgat4d* wild type and *Mgat4d*[−/−] germ cells prepared only 8 hr after males were treated at 43°C for 25 min, when no histological changes were apparent. Comparisons of DEGs and bioinformatics analyses using IPA, GSEA and EnrichR revealed that *Mgat4d*[−/−] heat-treated germ cells responded initially to heat stress, but did not sustain that response like wild type, heat-treated germ cells. Thus, *Mgat4d*[−/−] germ cells were less protected by autophagy or signaling pathways of inflammatory and proliferative responses. In addition, heat-treated *Mgat4d*[−/−] germ cells upregulated spermatogenic and spermiogenic genes to a greater extent than controls, indicative of the loss of a regulator of spermatogenesis - MGAT4D in this case. We previously showed that loss of MGAT1 in germ cells gave a similar upregulation of genes that promote spermatogenesis or spermiogenesis ^6^.

A key question for the future is to determine how MGAT4D protects against heat shock in male germ cells. Interestingly, *Mgat4d* transcripts are markedly reduced by the 43°C treatment and yet if MGAT4D is not present, germ cells are more sensitive to heat treatment, and if a *Mgat4d* transgene is present, germ cells are comparatively protected. Thus, the presence of MGAT4D, which may perdure in wild type germ cells after *Mgat4d* transcripts are reduced by heat stress, appears to facilitate the sustained heat stress response observed in wild type germ cells. How this is accomplished by a type II transmembrane Golgi glycoprotein may be related to the effects of Golgi glycosyltransferases on Golgi fragmentation. Some Golgi glycosyltransferases of the medial and trans Golgi compartments have been shown to facilitate Golgi fragmentation after heat shock ^33,34^. For example, the mucin O-glycan GlcNAcT CGNT3 promotes Golgi fragmentation following heat shock by interacting with myosin IIA via its cytoplasmic tail ^34^. MGAT4D is the most abundant protein in rat Golgi of male germ cells ^4^ and its loss after heat shock may protect the Golgi from fragmentation and protect Golgi glycosyltransferases and other Golgi residents, including molecules that protect from Inflammation and autophagy and that promote proliferation and survival, from degradation by the proteasome ^34^.

## Materials and Methods

### Mice

Mice carrying a conditional *Mgat4d* allele were generated from JM8A3.N1 ES cells carrying the targeting construct (Fig. 1) that were obtained from KOMP (project CSD79367). Targeted ES cells were injected into C57BL/6J blastocysts by the Gene Targeting Facility of the Albert Einstein College of Medicine. Chimeras were crossed to C57BL/6J mice and then to the FVB/NJ Stra8-iCre mice from Jackson Labs (Bar Harbor, Maine) Tg (Stra8-icre)1Reb/J (Stock no. 008208 | Stra8-iCre) to generate *Mgat4d* deleted mice carrying LacZ/Neo (*Mgat4d*-LacZ/Neo) or to Flp1-Cre mice B6.129S4-*Gt(ROSA)26Sor*^*tm1(FLP1)Dym*^/RainJ (Stock no. 009086 ROSA26:FLPe knock-in) to obtain mice carrying a conditional *Mgat4d* allele with *loxP* sites flanking exon 4 (*Mgat4d*[F/F]). The latter mice were crossed to mice carrying a Stra8-iCre transgene to generate conditional inactivation in spermatogonia to investigate spermatogenesis and fertility in males, or to generate mice with a whole body inactivation of *Mgat4d*. Transgenic mice used in this study were generated in Albert Einstein College of Medicine by the Transgenic Mouse Facility of the Albert Einstein College of Medicine on a C57BL/6J background. Two founders were characterized for each transgenic line. The constructs used are shown in Fig. 4. C57BL/6J and FVB/NJ mice were purchased from Jackson Laboratories (Stock No: 000664 and Stock No: 001800 respectively) and used for breeding. All mice carrying a transgene were kept as heterozygotes by crossing +/Tg with homozygote wild-type (+/+) mice. Mice were sacrificed by carbon dioxide asphyxiation followed by cervical dislocation. Testes were dissected free of surrounding tissue and weighed. Mouse experiments were performed following Albert Einstein College of Medicine Institutional Animal Care and Use Committee approved guidelines under the Institutional Animal Care and Use Committee (IACUC) protocol nos. 20080813, 20110803, 20140803 and 20170709.

### Antibodies

Anti-MGAT4D C-terminus pAb (Genemed, Torrance, CA) was obtained from a MGAT4D C-terminus peptide conjugate CGTQSSFPGREQHLKDNYY injected into rabbits. Anti-MGAT4D N-terminus pAb (Covance, Denver, PA; Genemed, Torrance, CA) was obtained with a MGAT4D-L N-terminal peptide conjugate GESVGDLRTVATAPWEGEQARGV injected into rabbits. Both pAbs were affinity purified on respective peptide columns. Anti-Myc mouse mAb 9E10 was from Covance (Denver, PA).

### Immunohistochemistry

Testes were fixed in Bouin’s fixative (#100503–962, Electron Microscopic Sciences, Radnor, PA) for 48 hr at room temperature (RT) then processed and paraffin-embedded by the Einstein Histology and Comparative Pathology Facility. Serial sections (5-6 μm) were collected on positively-charged slides. Immunohistochemistry was performed following the “IHC staining protocol for paraffin-embedded sections” from Abcam (http://www.abcam.com/protocols/). Briefly, testis sections were deparaffinized using Histo-Clear reagent Cat no. HS-200 (National Diagnostics, Atlanta, GA). We performed a heat-induced epitope retrieval with citrate buffer (10 mM sodium citrate, 0.05% Tween 20, pH 6.0) at 100°C for 20 min followed by 20 min period at room temperature in the same buffer. The tissue was permeabilized with 0.1% Triton X-100 in Tris-buffered saline (TBS) for 10 min and blocked for 1 hr at room temperature with 10% normal serum (from the same species as secondary antibody) and 1% BSA in TBS. The primary antibody was diluted in TBS with 1% BSA and incubated overnight at 4°C (unless otherwise indicated). Endogenous peroxidase was quenched by incubating slides in 1.5% H_2_O_2_ in TBS for 10 min and rinsed before incubation with the Biotinylated secondary antibody diluted in TBS containing 1% BSA, for 1 hr at room temperature. The samples were washed and Vectastain® ABC-HRP reagent (cat no. PK-6100, Vector laboratories, Inc. Burlingame, CA) was added and incubated at room temperature for 30 min. After rinsing, peroxidase substrate 3,3′diaminobenzidine (DAB) (Vector laboratories, Cat# SK-4100) was used to detect the antibody, following the manufacturer protocol. The tissue was counter-stained with Mayer’s Hematoxylin solution (cat no. MHS16-500ML, Sigma-Aldrich). The specimens were dehydrated with histo-clear and mounted using Permount® reagent (cat no. SP15- 100, Fisher Scientific, Fair Lawn, NJ). Testis section images were produced using 3DHistec Panoramic 250 Flash II slide scanner obtained with the Shared instrumentation Grant SIG# 1S10OD019961-01 to the Analytical Imaging Facility (AIF) of the Albert Einstein College of Medicine.

### Western-blot analysis

Testis tissue lysates were prepared using RIPA Lysis Buffer (cat no. 20-188, Millipore, Temecula, CA) and following the protocol “Preparation of lysate from tissues” from Abcam with modifications. Briefly, the testis tissue was homogenized in 1× RIPA, 01% SDS, 1× protease inhibitor cocktail (cat no. 05892791001, Roche Diagnostics GmbH, Mannheim, Germany) at a ratio of 0.5 ml buffer for 0.05 g of tissue. The lysate was incubated with constant agitation (orbital shaker) at 4°C for 2 hr and then centrifugated for 20 min at 12000 rpm at 4°C. The supernatant was transferred to a fresh tube and supplemented with 100% glycerol to a final concentration of 20% glycerol. Protein yield was measured using Bradford based colorimetric assay, (cat no. 500-0006, Bio-Rad Protein assay, Bio-Rad, Hercules, CA). Isolated germ cell were lysed in buffer containing 1% IGEPAL, 1%TX-100, 0.5% Deoxycholate and 1× protease inhibitor cocktail in water. Briefly, 100 µl of lysis buffer was used to homogenize 10^7^ cells. The lysate was incubated for 30 min on ice, then centrifugated 5 minutes at 5000 g. The supernatant was transferred to a fresh tube and supplemented with 100% glycerol to a final concentration of 20% glycerol. Protein levels were measured using the Bradford-based colorimetric assay. All samples were stored at −80°C.

### Apoptosis assay

Apoptosis induced DNA damage was measured using the ApopTag® Peroxidase *In Situ* Apoptosis Detection Kit (cat no. S7100, EMD Millipore, Temecula, CA) following the manufacturer’s protocol for paraffin-embedded tissue. Testis sections were deparaffinized using Histo-Clear reagent (cat no. HS-200, National Diagnostics, Atlanta, GA). Stained slides were scanned using a Perkin Elmer P250 high capacity slide scanner and images were analyzed using FIJI software to count foci ^35^.

### Germ cells isolation

Male germ cells were purified from testis following a modified protocol ^36–38^. Mice were sacrificed by CO_2_ asphyxiation followed by cervical dislocation and both testes were collected in 2 ml DMEM: F12 medium (cat no. 11330-032, Gibco, Grand Island, NY) on ice. The tunica albuginea was removed and tubules were transferred to 10 ml enzyme solution I (0.5 mg/ml collagenase Type I (cat no.C0130-1G, Sigma), 200 μg/ml DNase I (cat no. DN25-100 mg, Sigma) in F12 medium), briefly vortexed and incubated 30 min at 33°C in a shaking water bath (100 oscillations/min). Every 10 min an additional manual shaking was done to help tissue dissociation. The dispersed seminiferous tubules were allowed to sediment and the supernatant was discarded. Tubules were washed with 10 ml fresh F12 medium and resuspended in fresh F12 medium. The mixture was layered on 40 ml of 5% Percoll (cat no. 17-0891-02, GE Healthcare Bio-sciences AB, Uppsala, Sweden) in HBSS (cat no. 55-022-PB, Mediatech, Inc. Manassas, VA) and allowed to settle for 20 min at room temperature. The top 45 ml containing Leydig cells was discarded and the remaining 5 ml were transferred to a new tube containing 10 ml of enzyme solution II (200 μg/ml DNase I, 1 mg/ml trypsin (cat no.T4799-5G, Sigma-Aldrich, St Louis, MO) in F12 medium). The mixture was incubated for 40 min at 33°C in a shaking water bath (100 oscillations/min) and every 10 min, manual shaking. After tissue dissociation, 3 ml charcoal-stripped FBS were added and cells were resuspended using a 10 ml pipette to dissociate clumps. The suspension was filtered sequentially through a 70 μm (cat no. 352350, Falcon Corning Incorporated, Corning, NY) then 40 μm (cat no 352340) nylon cell strainer and centrifugated at 500 g for 10 min at 4°C. The cell pellet was resuspended in 1 ml PBS (calcium and magnesium free) and counted. Cells were stored as a dry pellet at −80°C and used for protein or RNA extraction.

### RNA isolation and RT-PCR

Testes or isolated germ cells were homogenized in TRIZOL reagent (cat no. 15596018, Invitrogen) following the manufacturer’s protocol for tissue or cell pellet, respectively. The isolated total RNA was dissolved in RNase-free water, an aliquot (2 μl) was used to measure nucleic acid concentration and the remainder was immediately stored at −80°C. Total RNA (3 μg) was used to synthesize cDNA (75 μl final volume) with the Verso cDNA Synthesis Kit (cat no. AB-1453/A, Appliedbiosystems, Thermo scientific Baltics UAB, Vilnius, Lithuania) following the manufacturer’s protocol. cDNA was tested for genomic DNA contamination using end-point PCR with *Actb* primers flanking an exon and intron sequence (Supplementary Table S6).

### Quantitative PCR (qRT-PCR)

cDNA obtained as described above was used to perform real time PCR. PowerUp™ SYBR™ Green Master Mix (cat no. A25742, Applied Biosystems, Thermo Scientific Baltics UAB, Vilnius, Lithuania) was mixed with each sample to a primer final concentration of 150 nm, following the manufacturer’s protocol and run on a master cycler (ViiA 7, Thermo Fisher). PCR conditions were 95°C for 30 sec, followed by 40 cycles at 95°C for 15 sec, 60°C for 15 sec and 72°C for 20 sec. Unless otherwise stated, gene expression relative to *Actb* and *Rps2* was calculated by the log2^ddCT^ method ^39^.

### Histological analysis

Hematoxylin and eosin (H&E) counter stained testis sections were analyzed by light microscopy (Zeiss Axiovert 200M, Göttingen, GERMANY) or scanned using a Perkin Elmer P250 high capacity slide scanner and processed using the proprietary software CaseViewer (3D Histech P250 high capacity slide scanner, Perkin Elmer, Waltham, MA).

### Mild heat stress treatment

This protocol was adapted from ^12,40,41^. Briefly, an adult male mouse was anaesthetized in an isoflurane chamber with a constant oxygen flow of 2 L/min and 3 % isoflurane for 1 min followed by 2.5 % isoflurane for 3 min. The mouse was quickly removed from the chamber and its nose was introduced into a nose cone with the same anaesthesia parameters for another 1 min. Testes were secured in the scrotum by manual massage and one third of the body (hind legs, tail and scrotum) was immersed in a 43°C or 33°C (control) water bath, supported by a plastic tube for 25 min. During the experiment, the isoflurane flow was reduced every 10 min by 0.5 % to reach 1.5 % at the end of the treatment (2.5 % for 5 min after introduction into the water bath, then 2 % for 10 min and followed by 1.5 % for another 10 min). After the heat treatment, mice were dried on paper towel, allowed to recover in a chamber with oxygen flow at 2 L/min and 0% isoflurane for 5 to 10 min, then returned to a cage to recover from the effects of anaesthesia on a heating pad. Testes and epididymis were harvested 8 hr or 24 hr after treatment.

### Microarray

Germ cell RNA (150 ng, RIN>7.9) was provided to the Genomics Core Facility of the Albert Einstein College of Medicine for conversion to cDNA, labeling and hybridization to a mouse Affymetrix Clariom™ D array previously known as GeneChip™ Mouse Transcriptome Array 1.0 (Affymetrix, Santa Clara, CA). Raw intensity data (.CEL files) were mapped to genes using custom CDF files (clariomdmousemmgencodegcdf from http://brainarray.mbni.med.umich.edu/Brainarray/Database/CustomCDF/genomic_curated_CDF.asp), and rma-normalized using the R/Bioconductor package affy ^42^. Differential gene expression was modeled using limma ^43^. Genes with Benjamini-Hochberg-adjusted p-values <0.05 and fold-change >1.5 or <−1.5 were defined as differentially expressed genes (DEGs).

### Gene set enrichment analysis and Ingenuity Pathway Analysis

Gene set enrichment analysis (GSEA) ^25,44^ was performed to determine enrichment of gene sets from the curated (C2), GO (C5), and oncogenic signatures (C6) and Hallmark collections. Gene list enrichment analysis was performed using EnrichR ^28^ and Ingenuity Pathway Analysis IPA (www.qiagen.com/ingenuity, QIAGEN, Redwood City, CA) for genes with fold-change ± 1.5, p < 0.05 and False discovery rate p < 0.05.

### Statistical analysis

The bar graphs in all figures represent the mean ± SEM. Unpaired, two-tailed Student’s t test or one way ANOVA was used to calculate *p*-value using Graph Pad Prism 7.0 (Graph Pad Software Inc., La Jolla, CA). Statistical significance was indicated by *p<0.05, **p<0.01, ***p<0.001 or ****p<0.0001.

## Acknowledgements

We thank Frederic Bard for helpful comments on the manuscript, and the Histopathology Core, the Flow Cytometry core, the Transgenic Mouse and Gene Targeting Facilities and the Analytical Imaging Core of the Albert Einstein College of Medicine, who performed core services supported by Albert Einstein Cancer Center grant P01 13330.

## Author contributions

AA performed all experiments on transgenic mice, all microarray and validation experiments, data curation, bioinformatics analyses, and co-wrote the manuscript; ML characterized the original cohort of *Mgat4d* control and KO mice and transgenic mice, performed heat shock experiments and analyses on the old cohort and edited the paper; BB performed bioinformatics analyses and interpretation and edited the paper; FB developed the *Mgat4d* conditional and global knockout mice, and the LacZ mice; JA performed and interpreted MALDI-IMS data; JP made transgenic constructs, and bred mutant and transgenic mice; SS characterized antibodies; PS conceived and directed experiments, curated and interpreted data and co-wrote the paper.

## Additional Information

### Data Availability

The data generated and/or analysed for the current study are available from the corresponding author on reasonable request. Microarray data are deposited in NCBI’s Gene Expression Omnibus (GEO) and are accessible through GEO serial accession number GSE137307.

### Funding

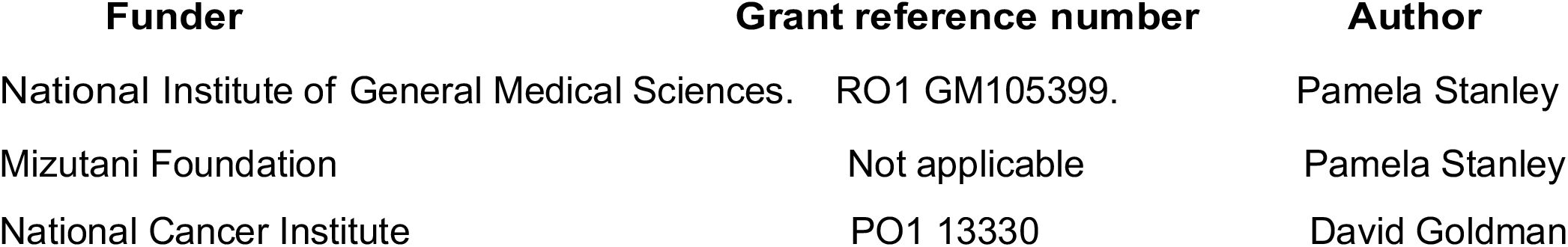

The funders had no role in study design, data collection or interpretation, or the decision to submit the work for publication.

