## supplemental information for "The Golgi Glycoprotein MGAT4D is an Intrinsic Protector of Testicular Germ Cells From Mild Heat Stress"

### Supplementary Methods

**MALDI-IMS.** Male mice were sacrificed, testes dissected, decapsulated, fixed in Bouin's fixative and embedded in paraffin. Sections of 6 µm were cut and mounted on Indium tin oxide (ITO) glass slides (Delta Technologies, Ltd., CO). Slides were deparaffinized and antigen retrieval was performed using citraconic anhydride buffer at pH 3 in a steamer for 30 min. Aqueous solution of PNGase F Prime (0.1 µg/µl, N-Zyme Scientifics, LLC, PA) was spray-coated on the slides using a TM Sprayer (HTX Technologies, LLC, NC) [1]. Slides were incubated at 37°C overnight in a humidified chamber and dried in a vacuum desiccator. Slides were then coated with α-cyano-4-hydroxycinnamic acid (CHCA) matrix (7 mg/ml in 50% Acetonitrile/ 0.1% TFA) using the TM Sprayer (HTX Technologies, LLC, NC). N-glycan imaging was performed by scanning and acquiring spectra from the entire tissue sections (m/z range 900-4,500) on an Ultraflextreme MALDI-TOF/TOF mass spectrometer equipped with a SmartBeam II laser at 1 kHz and 100 µm raster width. MALDI-TOF data were processed by FlexImaging software (v 4.1) to generate N-glycan ion maps.

### Supplementary figure legends

**Figure S1.** MALDI-IMS analysis of wild type and *Mgat4d*[-/-] testis sections. (**A**) Representative images of testis sections showing, in pseudo-color, the relative abundance of the predicted N-glycan shown on the left of each row. (**B**) MALDI-IMS intensity profiles of predicted N-glycans in *Mgat4d*[+/+] (top) and *Mgat4d*[-/-] (bottom) testis sections. (**C**) Graphical representation of the spectrum mean intensity ratio in testis sections from *Mgat4d*[-/-] (n=4 ) over *Mgat4d*[+/+] (n=3) for each N-glycan species. ***p<0.001 based on unpaired, two-tailed Student's t test assuming equal STDEV. (**D**) Graphical representation of the mean intensity ratio for testis sections from transgenic *Stra8-Mgat4d-Myc/Ldhc-Mgat4d-Myc* (n=3) or *Stra8-Mgat4d-Myc/Ldhc-Mgat4d-Myc/Prm1-Mgat4d-Myc* (n=2) transgenic mice over *Mgat4d*[+/+] (n=3) for each N-glycan species. N-glycans are represented using the Symbol Nomenclature for Glycans [2-4].

**Figure S2.** Peptide antibodies against MGAT4D. (**A**) Extracts from CHO transfectants stably-expressing Myc-tagged MGAT4D [5,6] were subjected to western blot analysis using pAbs raised in rabbits against the N-terminus unique to MGAT4D-L. (**B**) The same CHO extracts were subjected to western analysis using rabbit pAbs against a C-terminal peptide of MGAT4D (lanes 1,2). Myc-MGAT4D-L has a much stronger signal than MGAT4D-L-Myc, presumably because the Myc sequence perturbed the epitope. The pAb detects endogenous MGAT4D-L and MGAT4D-S in lane 4 in wild type germ cells, and those signals are absent from *Mgat4d*[-/-] germ cells in lane 6. Germ cell extracts from 28 dpp males expressing the *Stra8-Mgat4d-L-Myc* and *Ldhc-Mgat4d-L-Myc* transgenes (lane 8) or the *Ldhc-Mgat4d-L-Myc* transgene alone (lane 9) exhibited endogenous signals as expected, but no clear transgene signal.

**Figure S3.** Evidence of *Mgat4d* exon 4 excision. (**A**) Diagram representing exons in the mouse showing primers (A, B, C, D) used for PCR of germ cell cDNA. (**B**) Diagram extracted from TAC4 software representing *Mgat4d* exon transcript abundance in germ cell cDNA from *Mgat4d*[-/-] (KO) and *Mgat4d*[+/+] (WT). Exon 4 has a low signal in the KO preparation. Other exons are transcribed explaining why *Mgat4d* is represented in *Mgat4d*[-/-] microarray data. However, no protein was observed from these transcripts (Fig. 1).

**Figure S4.** (**A**) Top enriched Hallmark gene sets in *Mgat4d*[-/-] germ cells obtained from Gene Set Enrichment Analysis (GSEA) with Normalized Enrichment Scores (NES) as shown. Each gene set had a nominal *p* value <0.05 and FDR <25%. (**B**) Top enriched Hallmark gene sets in *Mgat4d*[+/+] germ cells with NES shown. Each gene set had a nominal *p* value <0.05 and FDR <25%. (**C**) Top section of the heat map of genes from the Hallmark spermatogenesis gene set enriched in *Mgat4d*[-/-] germ cells. (**D**) Top section of the heat map of genes from the Hallmark gene set TNF alpha signaling via NFκB enriched in *Mgat4d*[+/+] germ cells.

**Supplementary Table S1.** Genotyping Primers

| Gene or Transgene | Primer name | Sequence | Product length (bp) |
|---|---|---|---|
| *Mgat4d* floxed | FB77-Fw<br>FB78-Rev | TCCCACCCATGAAACAGTCT<br>GTACTGGAGCCCAAGCAGAA | 477 *Mgat4d*[+]<br>623 *Mgat4d*[F] |
| *Mgat4d* deleted | FB79-Fw<br>FB-80-Rev | CCAGAGCTTAGAAAGCTGGTGT<br>TCAGTAATGGCTTTAATGGTCTATTT | 1008 *Mgat4d*[+]<br>246 *Mgat4d*[-]<br>1152 *Mgat4d* [F] |
| *Stra8-iCre* | Stra-Fw | AGATGCCAGGACATCAGGAACCTG | 236 |
| *Stra8-Tg junction* | Stra-Rev | ATCAGCCACACCAGACACAGAGATC | 639 |
| *Ldhc-Tg junction* | Ldhc-*Mgat4d*-Fw<br>*Mgat4d*-Rev | TGGAAACCGTCTGGAGTCGT<br>TGGAGATGCAGAAACACGAGAAACC | 536 |
| *Prm1-Tg junction* | Prm1-*Mgat4d*-Fw<br>*Mgat4d*-Rev | TGAAGCACTTGATGGGGCCT<br>TGGAGATGCAGAAACACGAGAAACC | 857 |

**Supplementary Table S2.** Testis weights.

| **Mouse Background** | **Temp. (°C)** | **Genotype** | **Testis Weight (mg) (+/- SEM)** | **Testis/Body weight (+/- SEM)** |
|---|---|---|---|---|
| FVB | 33 | *Mgat4d*[-/-] (n=4) | 81.7 (1.2) | 0.002 (8.8E-05) |
| | 43 | *Mgat4d*[+/-] (n=2) | 71.5 (14.8) | 0.0017 (2.3E-04) |
| | | *Mgat4d*[-/-] (n=5) | 71.5 (3.9) | 0.0015 (4.0E-05) |
| C57Bl6/J | 33 | *Mgat4d*[+/+] (n=3) | 90.4 (3.6) | 0.0031 (9.4E-05) |
| | | Stra8-*Mgat4d-Myc* (n=3) | 99.3 (3.3) | 0.0031 (1.4E-04) |
| | | Ldhc-*Mgat4d-Myc* (n=4) | 96.8 (2.7) | 0.0026 (9.8E-05) |
| | | Prm1-*Mgat4d-Myc* (n=3) | 106.3 (2.7) | 0.0030 (7.4E-04) |
| | 43 | *Mgat4d*[+/+] (n=5) | 85.4 (1.1) | 0.0026 (3.2E-05) |
| | | Stra8-*Mgat4d-Myc* (n=11) | 86.6 (2) | 0.0025 (4.5E-05) |
| | | Ldhc-*Mgat4d-Myc* (n=7) | 88.1 (2) | 0.0024 (6.5E-05) |
| | | Prm1-*Mgat4d-Myc* (n=5) | 94.5 (3.6) | 0.0029 (1.6E-04) |

**Supplementary Table S3.** DEGs for *Mgat4d*[-/-] versus *Mgat4d*[+/+] treated at 33°C ($p<0.05$; $Log_2$ Fold change +/-0.585 ; FDR<0.05).

| Gene_id | REFSEQ | Gene Symbol | $Log_2$ Fold Change | P.Value | FDR P.Val |
|---|---|---|---|---|---|
| ENSMUSG00000035057.7 | NM_026233.2;NA | *Mgat4d* | -2.476254706 | 1.71199E-12 | 7.07687E-08 |
| ENSMUSG00000083857.1 | N/A | Gm12584 | 0.651408486 | 6.05467E-06 | 0.031285232 |
| ENSMUSG00000096243.1 | N/A | Gm24265 | 0.59577105 | 1.36822E-05 | 0.035348888 |
| ENSMUSG00000031919.6 | NM_025458.2 | *Tmed6* | 0.615438203 | 2.11808E-05 | 0.04457681 |

N/A, none available

**Supplementary Table S4** Top 20 Down-regulated genes in *Mgat4d*[-/-] vs *Mgat4d*[+/+] treated at 43°C ($p$<0.0; $Log_2$ Fold change +/-0.585; FDR<0.05).

| Gene_id | REFSEQ | Gene Symbol | $Log_2$ Fold Change | P.Value | FDR P.Val |
|---|---|---|---|---|---|
| ENSMUSG00000044734.16 | NM_025429.2;NA | *Serpinb1a* | -2.13291396 | 1.62393E-08 | 0.000466018 |
| ENSMUSG00000112023.1 | NM_001291892.1;NM_008147.2;NA; NM_001291893.1 | *Ly96* | -2.127378416 | 7.72306E-07 | 0.001596241 |
| ENSMUSG00000029561.17 | NM_011854.2;NA | *S100a11* | -2.095241568 | 5.88104E-05 | 0.009529246 |
| ENSMUSG00000107988.1 | NM_011485.5 | *Star* | -1.991773919 | 0.000273218 | 0.01981437 |
| ENSMUSG00000001173.15 | NM_177215.3;NA | *Gm5552* | -1.946398486 | 6.5394E-06 | 0.003790091 |
| ENSMUSG00000021843.17 | NM_008477.2;NA;NM_001293636.1; NM_001293635.1;NM_001347522.1 | *Osr2* | -1.886229424 | 7.99092E-06 | 0.004078031 |
| ENSMUSG00000024697.3 | NM_008137.4 | *Klk1b22* | -1.757911947 | 0.000205567 | 0.017063269 |
| ENSMUSG00000021025.8 | NM_010907.2;NA | *Gstm2-ps1* | -1.66489437 | 0.000239868 | 0.018637979 |
| ENSMUSG00000115207.1 | N/A | *Hspd1-ps5* | -1.649714912 | 0.000489712 | 0.025072341 |
| ENSMUSG00000058427.10 | NM_009140.2;NA | *Vmn2r-ps55* | -1.561583539 | 0.000323231 | 0.021174979 |
| ENSMUSG00000020120.15 | NA;NM_019549.2 | *Gm906* | -1.512253739 | 8.96726E-06 | 0.004310228 |
| ENSMUSG00000107737.1 | N/A | *AC113125.1* | -1.501234542 | 8.60716E-05 | 0.01161002 |
| ENSMUSG00000075538.2 | N/A | *Ly6e-ps1* | -1.471201428 | 4.64307E-05 | 0.008418001 |
| ENSMUSG00000102555.1 | N/A | *Gm47657* | -1.463936317 | 2.64999E-06 | 0.002625423 |
| ENSMUSG00000021109.13 | NM_001313919.1;NM_010431.2; NM_001313920.1;NA | *Cxcl1* | -1.432819232 | 2.97223E-06 | 0.002681799 |
| ENSMUSG00000032420.8 | NM_011851.4;NA | *Cd36* | -1.421730882 | 0.000385586 | 0.022681955 |
| ENSMUSG00000020044.13 | NM_011595.2;NA | *Gm25992* | -1.420700907 | 0.000153868 | 0.015183827 |
| ENSMUSG00000025779.10 | NM_016923.2;NM_001159711.1;NA | *Serpina3g* | -1.414715206 | 4.10519E-05 | 0.007713469 |
| ENSMUSG00000019850.11 | NM_009397.3;NM_001166402.1;NA | *Cyp11a1* | -1.403700796 | 0.001239342 | 0.04030738 |
| ENSMUSG00000028128.13 | NM_010171.3;NA | *Gm24613* | -1.38644583 | 0.001591758 | 0.045661687 |

N/A, none available

**Supplementary Table S5.** Top 20 Up-regulated genes for Mgat4d[-/-] vs Mgat4d[+/+] treated at 43°C ($p$<0.05; $Log_2$ Fold change +/-0.585; FDR<0.05).

| **Gene_id** | **REFSEQ** | **Gene Symbol** | **$Log_2$ Fold Change** | **P.Value** | **FDR P.Val** |
|---|---|---|---|---|---|
| ENSMUSG00000044734.16 | NM_025429.2;NA | *Gm26715* | 2.039467879 | 6.67287E-05 | 0.01021617 |
| ENSMUSG00000112023.1 | NM_001291892.1;NM_008147.2; NA;NM_001291893.1 | *Gm48565* | 1.977932812 | 0.000937941 | 0.034380536 |
| ENSMUSG00000029561.17 | NM_011854.2;NA | *Hspa1a* | 1.809603885 | 0.000293976 | 0.020389374 |
| ENSMUSG00000107988.1 | NM_010478.2 | *Hspa1b* | 1.767645726 | 0.000438609 | 0.024104592 |
| ENSMUSG00000001173.15 | NM_177215.3;NA | *Gm37692* | 1.325226203 | 9.74602E-05 | 0.012430179 |
| ENSMUSG00000021843.17 | NM_008477.2;NA;NM_001293636.1; NM_001293635.1;NM_001347522.1 | *Gm9176* | 1.159699663 | 0.000549294 | 0.026232849 |
| ENSMUSG00000024697.3 | NM_008137.4 | *4933414I15Rik* | 1.115823396 | 0.000122704 | 0.013705122 |
| ENSMUSG00000021025.8 | NM_010907.2;NA | *Gm5795* | 1.066409818 | 0.001758011 | 0.048433599 |
| ENSMUSG00000115207.1 | N/A | *Slc13a5* | 1.02736092 | 7.09263E-05 | 0.010437146 |
| ENSMUSG00000058427.10 | NM_009140.2;NA | *Gm5134* | 1.026790159 | 0.000453311 | 0.024424745 |
| ENSMUSG00000020120.15 | NA;NM_019549.2 | *Gdpd4* | 1.022429475 | 0.000182972 | 0.016250887 |
| ENSMUSG00000107737.1 | N/A | *Gm37432* | 1.01770496 | 5.5473E-05 | 0.009209182 |
| ENSMUSG00000075538.2 | N/A | *Catspere1* | 0.95199565 | 0.001811793 | 0.049246727 |
| ENSMUSG00000102555.1 | N/A | *Rnf144b* | 0.947574235 | 0.000356009 | 0.022238592 |
| ENSMUSG00000021109.13 | NM_001313919.1;NM_010431.2; NM_001313920.1;NA | *Gm27317* | 0.941739767 | 1.34066E-05 | 0.005263451 |
| ENSMUSG00000032420.8 | NM_011851.4;NA | *Gm19224* | 0.939180805 | 0.000708318 | 0.029695455 |
| ENSMUSG00000020044.13 | NM_011595.2;NA | *Gm26785* | 0.925613481 | 3.87817E-05 | 0.007526383 |
| ENSMUSG00000025779.10 | NM_016923.2;NM_001159711.1;NA | *Cdkl4* | 0.91217717 | 0.000921422 | 0.034121523 |
| ENSMUSG00000019850.11 | NM_009397.3;NM_001166402.1;NA | *n-R5s40* | 0.907045248 | 0.000270505 | 0.01981437 |
| ENSMUSG00000028128.13 | NM_010171.3;NA | *Sgpp2* | 0.904271954 | 0.000321063 | 0.021167112 |

N/A, none available

**Supplementary Table S6.** Quantitative RT-PCR Primers

| Gene | Gene ID | Primer name | cDNA (bp) | Sequence |
|---|---|---|---|---|
| *Mgat4d-L* | NM_026233.2<br>HM067443 | Long-Fw<br>Long-Rev | 129 | TGCCTGGGAGAAAGTGTTGGGGACC<br>CGTGGCAGCGTCACTGCCAACACCA |
| *Mgat4d-Myc* transgene | | Tr-Fw<br>Tr Rev | 124 | GGATTTCGAATTCAGTACAGACCAT<br>CAGATCTTCTTCAGAAATAAGTTTTTGTTC |
| *Star* | NM_011485.5 | Star-Fw<br>Star-Rev | 155 | TCCTCGCTACGTTCAAGCTG<br>ACGTCGAACTTGACCCATCC |
| *Osr2* | NM_054049.2 | Osr2-Fw<br>Osr2-Rev | 122 | ACATATGCAGACATCAAGCCCT<br>CCTGGGCTTCGCTAGAAGTT |
| *Serpinb1a* | NM_025429.2 | Serp-Fw<br>Serp-Rev | 154 | GGCTTTTGCATGACCTCCAG<br>GGCTTAAGGGTATCCACGCT |
| *Cyp11a1* | NM_019779.4 | Cyp11-Fw<br>Cyp11-Rev | 164 | GGTTTGGGGCAGAGACACTC<br>AGGTACCAGCTCCCTTTCCA |
| *Ly96* | NM_001159711.1 | Ly96-Fw<br>Ly96-Rev | 145 | TGCAACTCCTCCGATGCAAT<br>TACGCTTCGGCAACTTTGGA |
| *KlK1b22* | NM_010114.2 | Klk1b-Fw<br>KlK1b-Rev | 143 | TGTCCATCAAGCTCCATCCT<br>ACCATCACAGATCAGTGGGC |
| *Tmed6* | NM_025458.2 | Tmed6-Fw<br>Tmed6-Rev | 158 | TCACCTGCAGAAGAACCCAC<br>TCATTCCGATCAGCTCCACG |
| *Slc2a3* | NM_011401.4 | Slc2a-Fw<br>Slc2a-Rev | 174 | ACCACGAGGAGGATGTGGTAA<br>AATCTCTGCAAGGGGTGGAG |
| *Gdpd4* | NM_177696.3 | Gdpd-Fw<br>Gdpd-Rev | 149 | TGGTCGCCTAGGCTCATAGA<br>GCATTAGGGAGCAACCCACT |
| *Prss42* | NM_153099.1 | Prss-Fw<br>Prss-Rev | 325 | CCTCTTGCTCCTTCAGCCAA<br>AATACAATGGGCGGCAGTCA |
| *Pabpc6* | NM_001163836.1 | Pabp-Fw<br>Pabp-Rev | 194 | GAGCCGTAGGGCATACTGTG<br>ATGTGCTGGCAGTACGATGT |
| *Hspa1a* | NM_010479.2 | Hspa1a-Fw<br>Hspa1a-Rev | 183 | CGAGGAGGTGGATTAGAGGC<br>AGCCCACGTGCAATACACAA |

| | | | | |
|---|---|---|---|---|
| *Hspa1b* | NM_010478.2 | Hspa1b-Fw<br>Hspa1b-Rev | 176 | ATCAGTGGGCTGTACCAGGG<br>CCAAGCAGCTATCAAGTGCAA |
| *Crybg3* | NM_174848.3 | Cryb-Fw<br>Cryb-Rev | 134 | GGCTGCCCATCAGCTAGAAT<br>AAAACGGAATTCACGGCGTC |
| *Hsp90aa1* | NM_010480.5 | Hsp90-Fw<br>Hsp90-Rev | 103 | CGAAGCATAACGACGATGAGC<br>ACCTTTGTTCCACGACCCAT |
| *Egfr* | NM_007912.4 | Egfr-Fw<br>Egfr-Rev | 154 | GAAGTGTGGCCATCTGGGTA<br>CAGGGCAAGAGGGCAGAATC |
| *Hist1h2aa* | NM_175658.2 | Hist1-Fw<br>Hist1-Rev | 147 | GCCAAGGGAACTACGCACAA<br>GCAGGTGGCGAGGAGTAATG |
| *Selenop* | NM_001042613.2 | Selen-Fw<br>Selen-Rev | 152 | AACTCGTCAAAAGTCGTCCGT<br>CTATGTACCACTCCGGGGCT |
| *Dnaic2* | NM_001034878.3 | Dnaic-Fw<br>Dnaic-Rev | 172 | AAGACCTGGCAAAAAGAGGGAA<br>GAAGGCAAGGTGCTTGGAGG |
| *Bcl2l12* | NM_029410.3 | Bcl2l12-Fw<br>Bcl2l12-Rev | 124 | TCTTCTCTAGCCGGGAAAGC<br>CGGCTCAATTCCATGGCTAGT |
| *Degs1* | NM_007853.5 | Degs1-Fw<br>Degs1-Rev | 116 | CGAGAGGAGTTCGAATGGGTC<br>CAGATCAGGTTGTGGTCAGGT |
| *Dmrt1* | NM_015826.5 | Dmrt1-Fw<br>Dmrt1-Rev | 129 | TACTCAGAAGCCAAAGCCAGT<br>GGACGCAGACTCACATTCCAG |
| *Socs3* | NM_007707.3 | Socs3-Fw<br>Socs3-Rev | 110 | CAAGGCCGGAGATTTCGCTT<br>GGAGCCAGCGTGGATCTG |
| *Rps2* | NM_008503.5 | Rps2-Fwr<br>Rps2-Rev | 112 | CTGACTCCCGACCTCTGGAAA<br>GAGCCTGGGTCCTCTGAACA |
| *Actb* | NM_007393.5 | b-Act-Fwr<br>b-Act-Rev | 195 | GGCTCCTAGCACCATGAAGAT<br>TAAAACGCAGCTCAGTAACAGTC |

**Supplementary Table S7.** Top upstream regulators at 43ºC sorted by p-value of overlap.

| Upstream Regulator | Expr Log Ratio | Molecule Type | Predicted State | Activation z-score | Flags | p-value of overlap | Mechanistic Network |
|---|---|---|---|---|---|---|---|
| lipopolysaccharide | | chemical drug | Inhibited | -5.4 | bias | 1.5E-20 | 114 (15) |
| dexamethasone | | chemical drug | | -1.8 | | 8.2E-20 | 136 (17) |
| TGFB1 | -0.507 | growth factor | Inhibited | -3.3 | | 3.8E-19 | 126 (17) |
| TNF | -0.379 | cytokine | Inhibited | -3.8 | | 3.9E-19 | 126 (17) |
| IFNG | | cytokine | Inhibited | -4.3 | | 1.0E-18 | 124 (17) |
| IL4 | | cytokine | | -1.9 | bias | 6.0E-18 | 114 (15) |
| IL1B | | cytokine | Inhibited | -3.9 | | 5.1E-17 | 125 (14) |
| dihydrotestosterone | | chemical - endogenous mammalian | Inhibited | -2.2 | bias | 2.5E-16 | 124 (20) |
| forskolin | | chemical toxicant | Inhibited | -3.7 | bias | 3.7E-16 | 133 (24) |
| tetradecanoylphorbol acetate | | chemical drug | Inhibited | -5.3 | bias | 4.7E-16 | 144 (22) |
| AGT | -1.091 | growth factor | Inhibited | -3.8 | bias | 8.8E-15 | 118 (17) |
| SB203580 | | chemical - kinase inhibitor | Activated | 2.9 | bias | 1.3E-14 | 123 (18) |
| cigarette smoke | | chemical toxicant | Inhibited | -3.0 | bias | 2.0E-14 | 118 (22) |
| tretinoin | | chemical - endogenous mammalian | Inhibited | -3.8 | | 2.6E-14 | 133 (19) |
| NFKBIA | -1.104 | transcription regulator | | -1.5 | | 2.9E-14 | 118 (17) |
| STAT6 | | transcription regulator | | 0.6 | | 5.0E-14 | 119 (15) |
| NR3C1 | | ligand-dependent nuclear receptor | | 0.9 | | 8.1E-14 | 141 (20) |
| NFE2L2 | | transcription regulator | Inhibited | -2.8 | bias | 9.9E-14 | 89 (15) |
| beta-estradiol | | chemical - endogenous mammalian | | -1.9 | | 1.9E-13 | 115 (15) |
| EGF | | growth factor | Inhibited | -3.2 | bias | 2.5E-13 | 131 (22) |

Top upstream regulators at 43ºC sorted by Expression Log Ratio.

| Upstream Regulator | Expr Log Ratio | Molecule Type | Predicted Activation State | Activation z-score | Flags | p-value of overlap | Mechanistic Network |
|---|---|---|---|---|---|---|---|
| LY96 | -2.127 | transmembrane receptor | | | | 7.3E-04 | 108 (20) |
| STAR | -1.992 | transporter | | | | 7.9E-03 | |
| OSR2 | -1.886 | transcription regulator | | | | 3.4E-02 | |
| CXCL2 | -1.433 | cytokine | | -1.1 | bias | 2.7E-05 | 79 (13) |
| ANXA5 | -1.332 | transporter | | | | 7.6E-04 | 46 (7) |
| IL1A | -1.313 | cytokine | Inhibited | -2.7 | bias | 1.7E-09 | 91 (12) |
| F3 | -1.229 | transmembrane receptor | | | | 4.3E-02 | |
| NFKBIA | -1.104 | transcription regulator | | -1.5 | | 2.9E-14 | 118 (17) |
| AGT | -1.091 | growth factor | Inhibited | -3.8 | bias | 8.8E-15 | 118 (17) |
| FXYD1 | -1.081 | ion channel | | | | 1.1E-02 | |
| HPGD | -1.07 | enzyme | | | | 2.3E-02 | |
| SLC13A5 | 1.027 | transporter | | | | 4.5E-02 | |
| HSPA1A/HSPA1B | 1.81 | enzyme | | | | 3.8E-04 | 64 (7) |

**Supplementary Table S8.** Top Networks in *Mgat4d*[-/-] versus *Mgat4d*[+/+] at 43°C.

| Rank | Score | Focus Molecules | Top Diseases and Functions |
|---|---|---|---|
| 1 | 52 | 28 | [DNA Replication, Recombination, and Repair, Nucleic Acid Metabolism, Small Molecule Biochemistry] |
| 2 | 30 | 19 | [Cell Death and Survival, Cellular Compromise, Endocrine System Development and Function] |
| 3 | 26 | 17 | [Lipid Metabolism, Molecular Transport, Small Molecule Biochemistry] |
| 4 | 26 | 17 | [Cell-To-Cell Signaling and Interaction, Cellular Movement, Tissue Development] |
| 5 | 24 | 16 | [Carbohydrate Metabolism, Cellular Compromise, Small Molecule Biochemistry] |
| 6 | 22 | 15 | [Cardiovascular Disease, Drug Metabolism, Glutathione Depletion In Liver] |
| 7 | 20 | 14 | [Cardiovascular System Development and Function, Dermatological Diseases and Conditions, Organismal Injury and Abnormalities] |
| 8 | 20 | 14 | [Amino Acid Metabolism, Cancer, Small Molecule Biochemistry] |
| 9 | 18 | 13 | [Cancer, Gastrointestinal Disease, Organismal Injury and Abnormalities] |
| 10 | 18 | 13 | [Gene Expression, RNA Damage and Repair, RNA Post-Transcriptional Modification] |
| 11 | 18 | 13 | [Cellular Compromise, Cellular Development, Cellular Growth and Proliferation] |
| 12 | 18 | 13 | [Cell Cycle, Connective Tissue Development and Function, Tissue Morphology] |
| 13 | 16 | 12 | [Cardiac Dilation, Ophthalmic Disease, Organismal Injury and Abnormalities] |
| 14 | 16 | 12 | [Hematological System Development and Function, Inflammatory Response, Tissue Morphology] |
| 15 | 14 | 11 | [Cellular Development, Cellular Growth and Proliferation, Hematological System Development and Function] |
| 16 | 14 | 11 | [Metabolic Disease, Neurological Disease, Organismal Injury and Abnormalities] |
| 17 | 14 | 11 | [Cell Cycle, Lipid Metabolism, Small Molecule Biochemistry] |
| 18 | 11 | 9 | [Hereditary Disorder, Neurological Disease, Organismal Injury and Abnormalities] |
| 19 | 2 | 1 | [Cell Cycle, Cell Morphology, Cellular Movement] |
| 20 | 2 | 1 | [Cellular Compromise, Inflammatory Response, Protein Degradation] |
| 21 | 2 | 1 | [Connective Tissue Development and Function, Organismal Injury and Abnormalities, Reproductive System Disease] |

**Supplementary Table S9.** Top most represented diseases and biofunctions in *Mgat4d*[-/-] vs C57BL/6J *Mgat4d*[+/+] germ cells treated at 43°C. Sorted by activation Z-score.

| Categories | Diseases or Functions | *p*-value | Predicted State | Activation z-score |
|---|---|---|---|---|
| Organismal Survival | Morbidity or mortality | 1.62E-12 | Increased | 6.137 |
| Organismal Survival | Organismal death | 2.04E-12 | Increased | 6.01 |
| Nutritional Disease,Organismal Injury and Abnormalities | Cachexia | 6.88E-6 | Increased | 2.773 |
| Inflammatory Response | Inflammation of body cavity | 2.10E-15 | Increased | 2.679 |
| Inflammatory Disease,Inflammatory Response,Organismal Injury and Abnormalities,Respiratory Disease | Inflammation of lung | 1.14E-08 | Increased | 2.633 |
| Infectious Diseases | Infection of mammalia | 4.61E-7 | Increased | 2.576 |
| Inflammatory Response,Respiratory Disease | Inflammation of respiratory system component | 1.97E-12 | Increased | 2.549 |
| Cardiovascular Disease,Organismal Injury and Abnormalities | Cardiac lesion | 4.24E-6 | Increased | 2.547 |
| Inflammatory Response,Organismal Injury and Abnormalities | Inflammation of organ | 1.17E-16 | Increased | 2.529 |
| Infectious Diseases | Systemic inflammatory response syndrome and/or sepsis | 7.11E-08 | Increased | 2.414 |
| Endocrine System Development and Function,Lipid Metabolism,Small Molecule Biochemistry,Vitamin and Mineral Metabolism | Steroidogenesis of hormone | 9.86E-09 | Decreased | -3.477 |
| Lipid Metabolism,Small Molecule Biochemistry,Vitamin and Mineral Metabolism | Metabolism of terpenoid | 1.37E-10 | Decreased | -3.543 |
| Cellular Movement | Invasion of cells | 7.74E-08 | Decreased | -3.546 |
| Endocrine System Development and Function,Small Molecule Biochemistry | Synthesis of hormone | 3.99E-08 | Decreased | -3.633 |
| Endocrine System Development and Function,Small Molecule Biochemistry | Metabolism of hormone | 1.51E-7 | Decreased | -3.633 |
| Cellular Movement | Cell movement of tumor cell lines | 4.27E-7 | Decreased | -3.721 |
| Organismal Development | Size of body | 5.18E-09 | Decreased | -3.937 |
| Lipid Metabolism,Small Molecule Biochemistry,Vitamin and Mineral Metabolism | Synthesis of steroid | 1.48E-09 | Decreased | -4.209 |
| Lipid Metabolism,Small Molecule Biochemistry | Synthesis of lipid | 3.45E-12 | Decreased | -4.255 |
| Lipid Metabolism,Small Molecule Biochemistry,Vitamin and Mineral Metabolism | Synthesis of terpenoid | 2.55E-10 | Decreased | -4.426 |

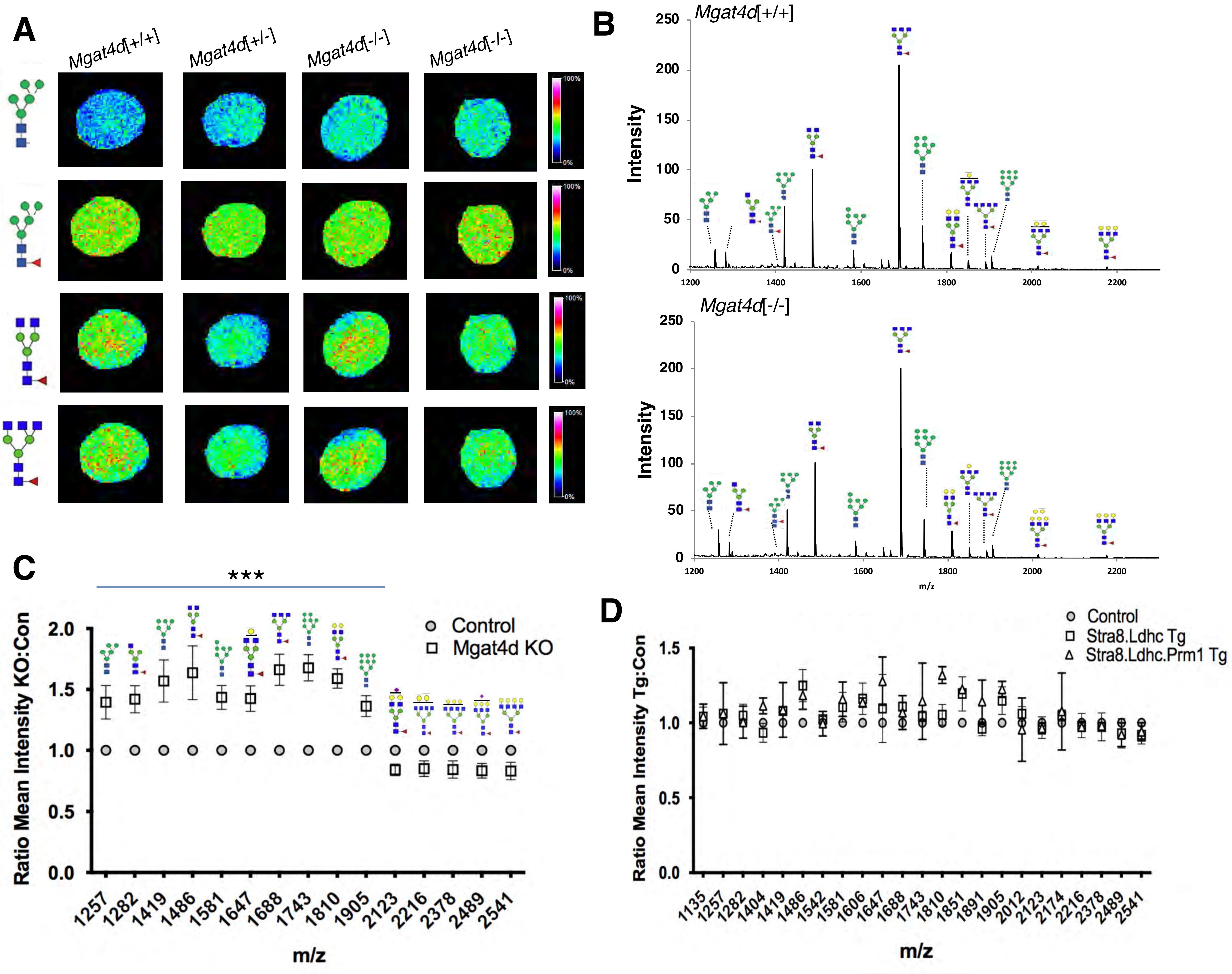
A
Mgat4d[+/+]
Mgat4d[+/-]
Mgat4d[-/-]
Mgat4d[-/-]
100%
0%
B
Mgat4d[+/+]
Mgat4d[-/-]
Intensity
m/z
C
***
Control
Mgat4d KO
Ratio Mean Intensity KO:Con
m/z
D
Control
Stra8.Ldhc Tg
Stra8.Ldhc.Prm1 Tg
Ratio Mean Intensity Tg:Con
m/z

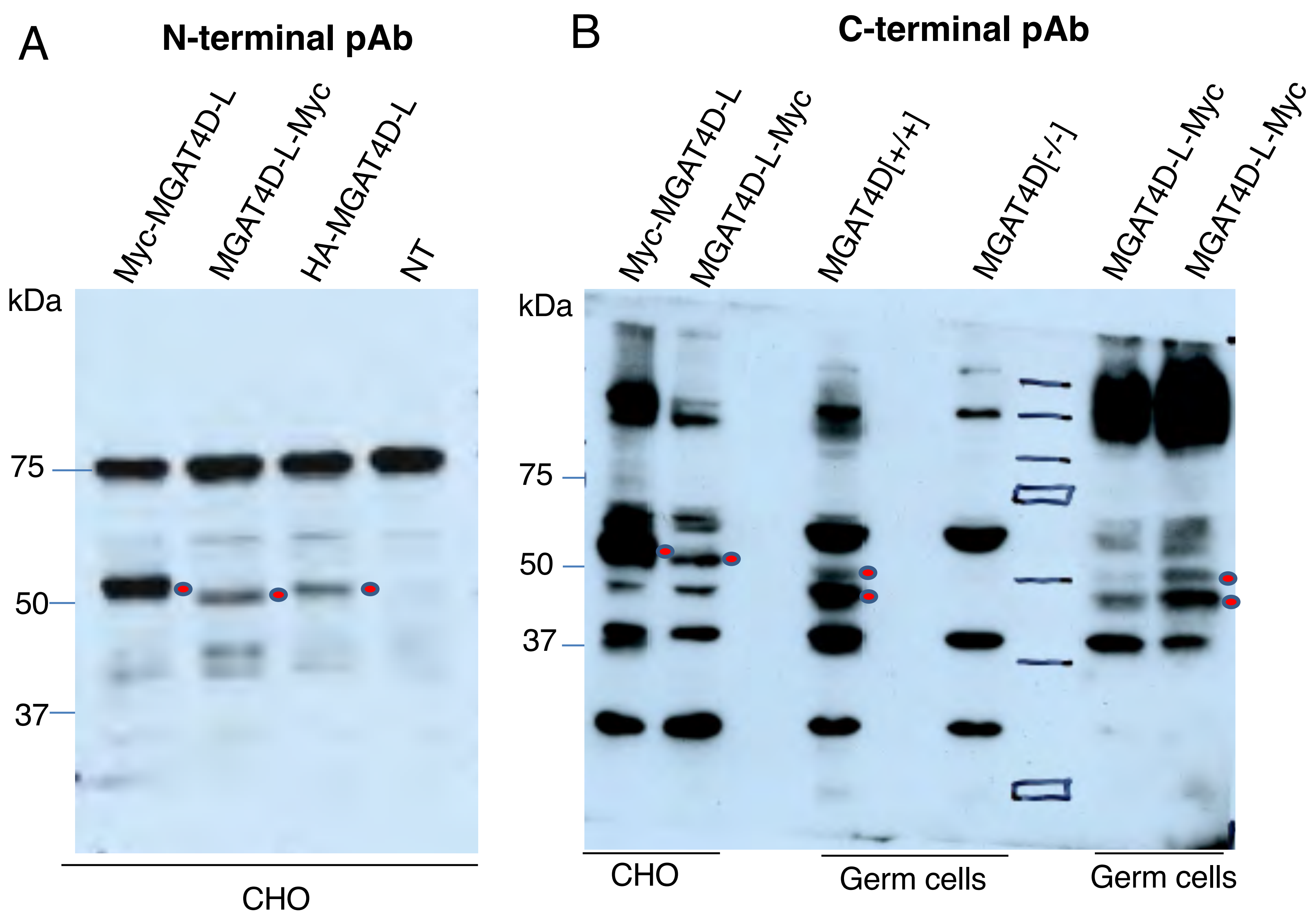
A
N-terminal pAb
Myc-MGAT4D-L
MGAT4D-L-Myc
HA-MGAT4D-L
NT
kDa
75
50
37
CHO
B
C-terminal pAb
Myc-MGAT4D-L
MGAT4D-L-Myc
MGAT4D[+/+]
MGAT4D[-/-]
MGAT4D-L-Myc
MGAT4D-L-Myc
kDa
75
50
37
CHO
Germ cells
Germ cells

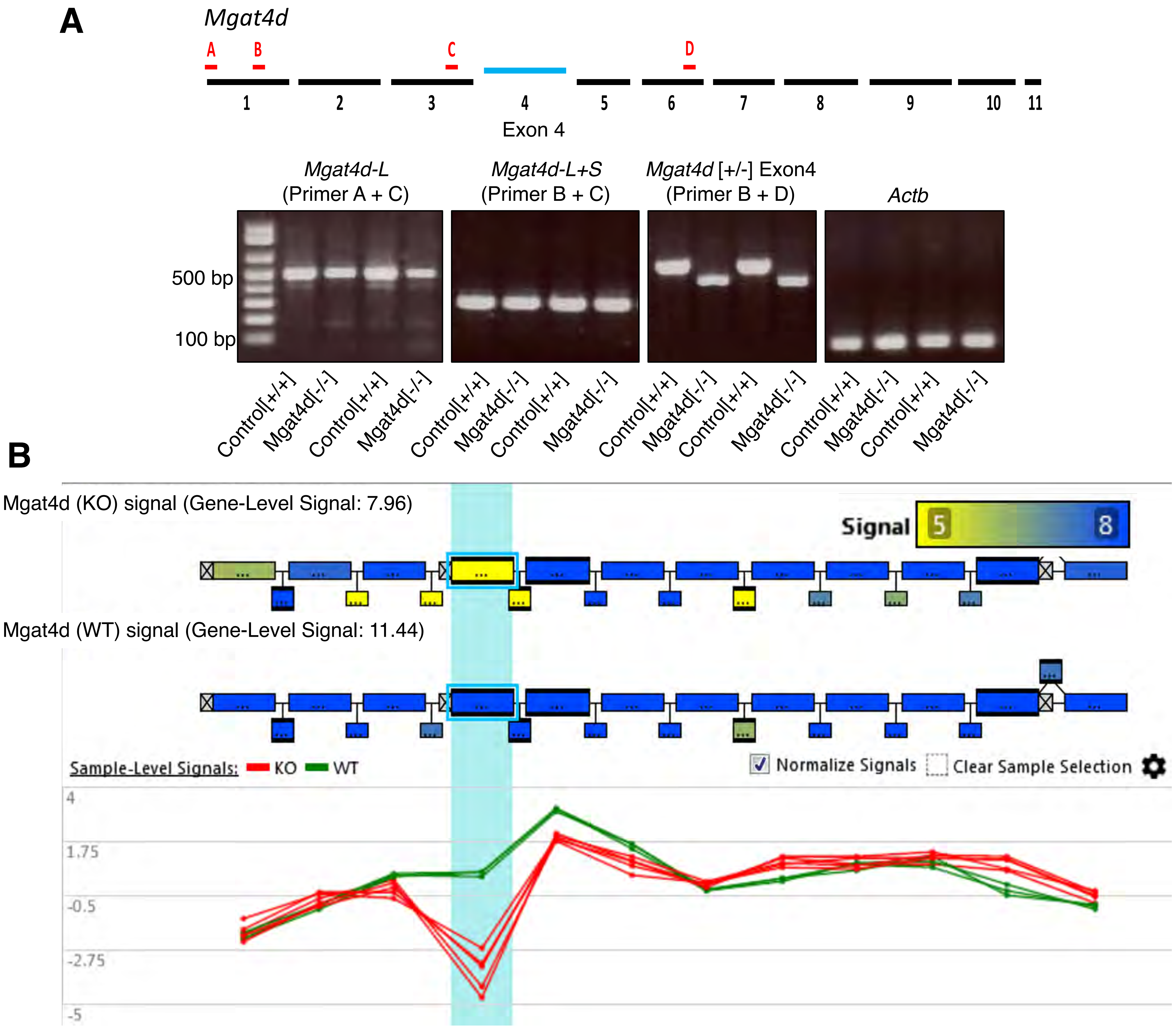
A
Mgat4d
A
B
C
D
1
2
3
4
5
6
7
8
9
10
11
Exon 4
Mgat4d-L
(Primer A + C)
Mgat4d-L+S
(Primer B + C)
Mgat4d [+/-] Exon4
(Primer B + D)
Actb
500 bp
100 bp
Control[+/+]
Mgat4d[-/-]
Control[+/+]
Mgat4d[-/-]
Control[+/+]
Mgat4d[-/-]
Control[+/+]
Mgat4d[-/-]
Control[+/+]
Mgat4d[-/-]
Control[+/+]
Mgat4d[-/-]
Control[+/+]
Mgat4d[-/-]
Control[+/+]
Mgat4d[-/-]
B
Mgat4d (KO) signal (Gene-Level Signal: 7.96)
Signal
5
8
Mgat4d (WT) signal (Gene-Level Signal: 11.44)
Sample-Level Signals: KO WT
Normalize Signals
Clear Sample Selection
4
1.75
-0.5
-2.75
-5

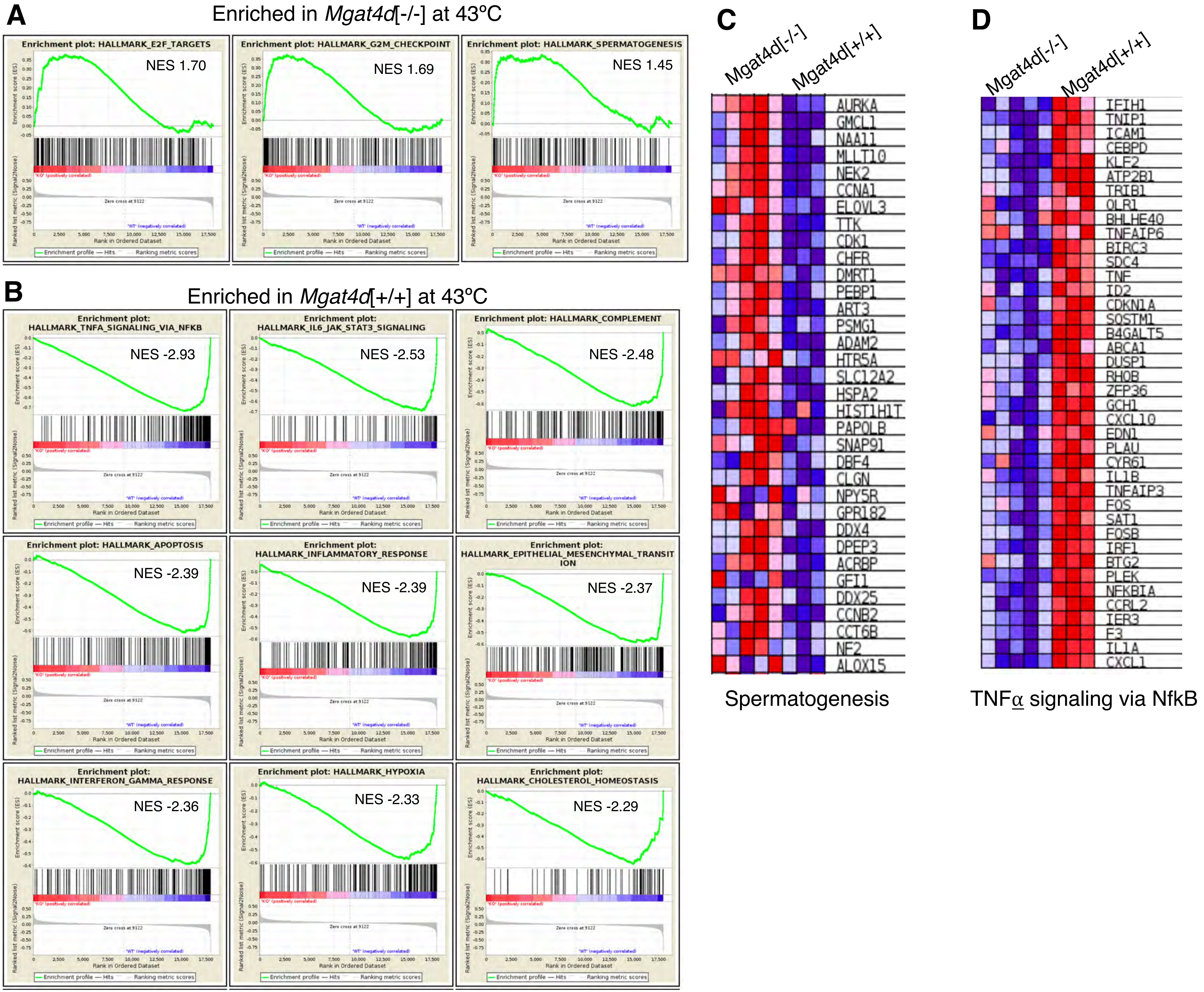
A
Enriched in Mgat4d[-/-] at 43°C
Enrichment plot: HALLMARK_E2F_TARGETS
NES 1.70
Enrichment plot: HALLMARK_G2M_CHECKPOINT
NES 1.69
Enrichment plot: HALLMARK_SPERMATOGENESIS
NES 1.45
B
Enriched in Mgat4d[+/+] at 43°C
Enrichment plot: HALLMARK_TNFA_SIGNALING_VIA_NFKB
NES -2.93
Enrichment plot: HALLMARK_IL6_JAK_STAT3_SIGNALING
NES -2.53
Enrichment plot: HALLMARK_COMPLEMENT
NES -2.48
Enrichment plot: HALLMARK_APOPTOSIS
NES -2.39
Enrichment plot: HALLMARK_INFLAMMATORY_RESPONSE
NES -2.39
Enrichment plot: HALLMARK_EPITHELIAL_MESENCHYMAL_TRANSITION
NES -2.37
Enrichment plot: HALLMARK_INTERFERON_GAMMA_RESPONSE
NES -2.36
Enrichment plot: HALLMARK_HYPOXIA
NES -2.33
Enrichment plot: HALLMARK_CHOLESTEROL_HOMEOSTASIS
NES -2.29
C
Mgat4d[-/-]
Mgat4d[+/+]
AURKA
GMCL1
NAA11
MLLT10
NEK2
CCNA1
ELOVL3
TTK
CDK1
CHFR
DMRT1
PEBP1
ART3
PSMG1
ADAM2
HTR5A
SLC12A2
HSPA2
HIST1H1T
PAPOLB
SNAP91
DBF4
CLGN
NPY5R
GPR182
DDX4
DPEP3
ACRBP
GFI1
DDX25
CCNB2
CCT6B
NF2
ALOX15
Spermatogenesis
D
Mgat4d[-/-]
Mgat4d[+/+]
IFIH1
TNIP1
ICAM1
CEBPD
KLF2
ATP2B1
TRIB1
OLR1
BHLHE40
TNFAIP6
BIRC3
SDC4
TNF
ID2
CDKN1A
SQSTM1
B4GALT5
ABCA1
DUSP1
RHOB
ZFP36
GCH1
CXCL10
EDN1
PLAU
CYR61
IL1B
TNFAIP3
FOS
SAT1
FOSB
IRF1
BTG2
PLEK
NFKBIA
CCRL2
IER3
F3
IL1A
CXCL1
TNFα signaling via NfkB
